# Sparse and stereotyped encoding implicates a core glomerulus for ant alarm behavior

**DOI:** 10.1101/2022.12.29.522224

**Authors:** Taylor Hart, Dominic Frank, Lindsey E. Lopes, Leonora Olivos-Cisneros, Kip D. Lacy, Waring Trible, Amelia Ritger, Stephany Valdés-Rodríguez, Daniel J. C. Kronauer

## Abstract

Ants communicate via large arrays of pheromones and possess expanded, highly complex olfactory systems, with antennal lobes in the brain comprising ~500 glomeruli. This expansion implies that odors could activate hundreds of glomeruli, which would pose challenges for higher order processing. To study this problem, we generated the first transgenic ants, expressing the genetically encoded calcium indicator GCaMP6s in olfactory sensory neurons. Using two-photon imaging, we mapped complete glomerular responses to four ant alarm pheromones. Alarm pheromones robustly activated ≤6 glomeruli, and activity maps for the three pheromones inducing panic-alarm in our study species converged on a single glomerulus. These results demonstrate that, rather than using broadly tuned combinatorial encoding, ants employ precise, narrowly tuned, and stereotyped representation of alarm pheromone cues. The identification of a central sensory hub glomerulus for alarm behavior suggests that a simple neural architecture is sufficient to translate pheromone perception into behavioral outputs.

## Introduction

Eusocial insects, like ants and honeybees, use vast arrays of pheromones to communicate information with conspecifics and to regulate colony life. These adaptations correspond to elaborations of the chemosensory system, which are particularly striking in ants. Insect olfactory systems have a conserved organization, with olfactory sensory neurons (OSNs) in peripheral sensory organs innervating glomeruli in the antennal lobes (ALs) in the brain (Strausfeld and Hildebrand, 1999; Vosshall et al., 2000; Zhao and McBride, 2020). Much of the detailed knowledge of insect olfactory system development, anatomy, and neural function comes from studies of the vinegar fly *Drosophila melanogaster*. However, ants possess an order of magnitude more odorant receptor genes (ORs) and AL glomeruli than *Drosophila* (Mysore et al., 2009; Kelber et al., 2010; Smith et al., 2011; Zhou et al., 2012; Zhou et al., 2015; McKenzie et al., 2016; McKenzie and Kronauer, 2018; Ryba et al., 2020; Trible et al., 2017; Ferguson et al., 2021; Benton 2022). In *Drosophila*, the ~50 AL glomeruli each receive input from a functional class of OSNs and have stereotyped positions across individuals, which allowed the creation of atlases mapping odor-evoked response functions for each glomerulus (Stocker et al., 1990; Stocker, 1994; Gao et al., 2000; Vosshall et al., 2000; Wang et al., 2003). By contrast, little is known about how odors are represented in the more complex olfactory system of ants with its ~500 AL glomeruli.

Here we focus on the neural representation of alarm pheromones, “danger” signals that are chemically well characterized across several ant species. Stimulating individuals with volatile alarm pheromones is experimentally simple and quickly elicits behavioral responses, which makes these pheromones attractive models for studying the neurobiological basis of chemical communication. Upon perception of the pheromone, locomotion usually increases, and aggression or “panic” commences (Wilson and Regnier, 1971). The alarm response can culminate in nest evacuation, where ants leave the nest carrying brood (Duffield et al., 1976; Smith and Haight, 2008). Specific features of alarm behavior vary with context, species, and specific mixtures and concentrations of chemicals, but frequently include either frenzied panic responses or attraction to the alarm source, as well as changes in the posture of antennae, mandibles, and the sting (Blum, 1969; Vander Meer and Alonso, 1998).

Alarm pheromone representation has been investigated using calcium dyes to record activity from subsections of the AL in several carpenter ant species (Galizia et al., 1999; Zube et al., 2008; Brandstaetter et al., 2011) and honeybees (Joerges et al., 1997; Galizia et al., 1998; Sachse et al., 1999; Guerrieri et al., 2005; Haase et al., 2011; Carcaud et al., 2015; Paoli and Galizia, 2021). These studies found broad, multi-glomerular activation patterns without evidence for specialized glomerulus clusters, similar to the combinatorial representation of general odorants in *Drosophila* (Joerges et al., 1997; Laurent, 1999; Wang et al., 2003; Hallem and Carlson, 2006; Carcaud et al., 2015; Münch and Galizia, 2016).

Such a combinatorial model with broad tuning implies that odor mixtures could potentially activate combinations of hundreds of glomeruli in the expanded ant AL. Because the number of potential combinations of glomeruli grows super-linearly with each additional glomerulus, this scenario poses much bigger challenges for higher order neurons in ants vs. *Drosophila* with respect to decoding multicomponent olfactory signals, detecting and identifying pheromones, and activating appropriate behavioral responses. In contrast, narrower tuning, where most odorants only activate a small number of glomeruli, could simplify the neural architecture necessary for processing odor information in the complex olfactory environment of an ant colony and ensure that pheromone signals can be rapidly and accurately perceived. Consistent with this alternative model is the relatively narrow tuning observed for at least some ant ORs (Pask et al., 2017; Slone et al., 2017).

The ant olfactory system also differs from that of *Drosophila* in several developmental properties that might be linked to its increased complexity (Trible et al., 2017; Yan et al., 2017; Duan and Volkan, 2020; Ryba et al., 2020). Based on these differences, it has been suggested that ants, similar to mice but unlike flies, might rely on intrinsic features of ORs for OSN axon guidance and AL patterning (Duan and Volkan, 2020; Ryba et al., 2020). This in turn could translate to increased developmental plasticity in the olfactory system. In both mice and *Drosophila*, olfactory glomeruli receiving input from a defined class of OSNs are consistently located in the same anatomical region, but at the local scale, homologous mouse glomeruli vary substantially in their spatial location across individuals, and even across the left/right axis within a single individual (Strotmann et al., 2000; Schaefer et al., 2001; Lodovichi and Belluscio, 2012; Zapiec and Mombaerts, 2015). Whether the level of anatomical-functional stereotypy of the ant olfactory glomeruli more closely resembles *Drosophila* or mice has not been assessed. However, the number of glomeruli in ants varies with sex, caste, and worker body size (Mysore et al., 2009; Kelber et al., 2010; Kuebler et al., 2010; McKenzie et al., 2016), suggesting that stereotypy may be low.

We studied the representation of alarm pheromones in the clonal raider ant *Ooceraea biroi*, an experimentally tractable species that lives in small colonies, reproduces asexually, and preys on other ants (Oxley et al., 2014; Trible et al., 2017; Chandra et al., 2021). We implemented the first neurogenetic tools in ants by developing a piggyBac transgenesis protocol to generate lines that express the genetically encoded calcium indicator GCaMP6s in OSNs. We then examined the relationships between behavioral outputs of alarm pheromone stimuli and single glomerulus-resolution, whole-AL calcium responses for four ant alarm pheromones.

## Results

### Alarm pheromones elicit a range of behavioral responses

The alarm pheromones 4-methyl-3-heptanone and 4-methyl-3-heptanol have previously been extracted from clonal raider ants and verified to elicit panic alarm responses in a colony bioassay, both alone and as a 9:1 blend that mimics the relative abundance of these compounds in ant head extracts (Fig. 1A-B, Table S1; Lopes et al., 2022). Two chemically related compounds, 4-methyl-3-hexanol and 6-methyl-5-hepten-2-one, act as alarm pheromones in other ant species but were not found in clonal raider ant chemical extracts (Fig. 1A, Table S1; Bernardi et al., 1967; McGurk, 1968; Duffield et al., 1977; Pasteels et al., 1980; Pasteels et al., 1981; Morgan et al., 1992; Keegans et al., 1993; Oldham et al., 1994; Han et al., 2022; Lopes et al. 2022).

**Figure 1.**
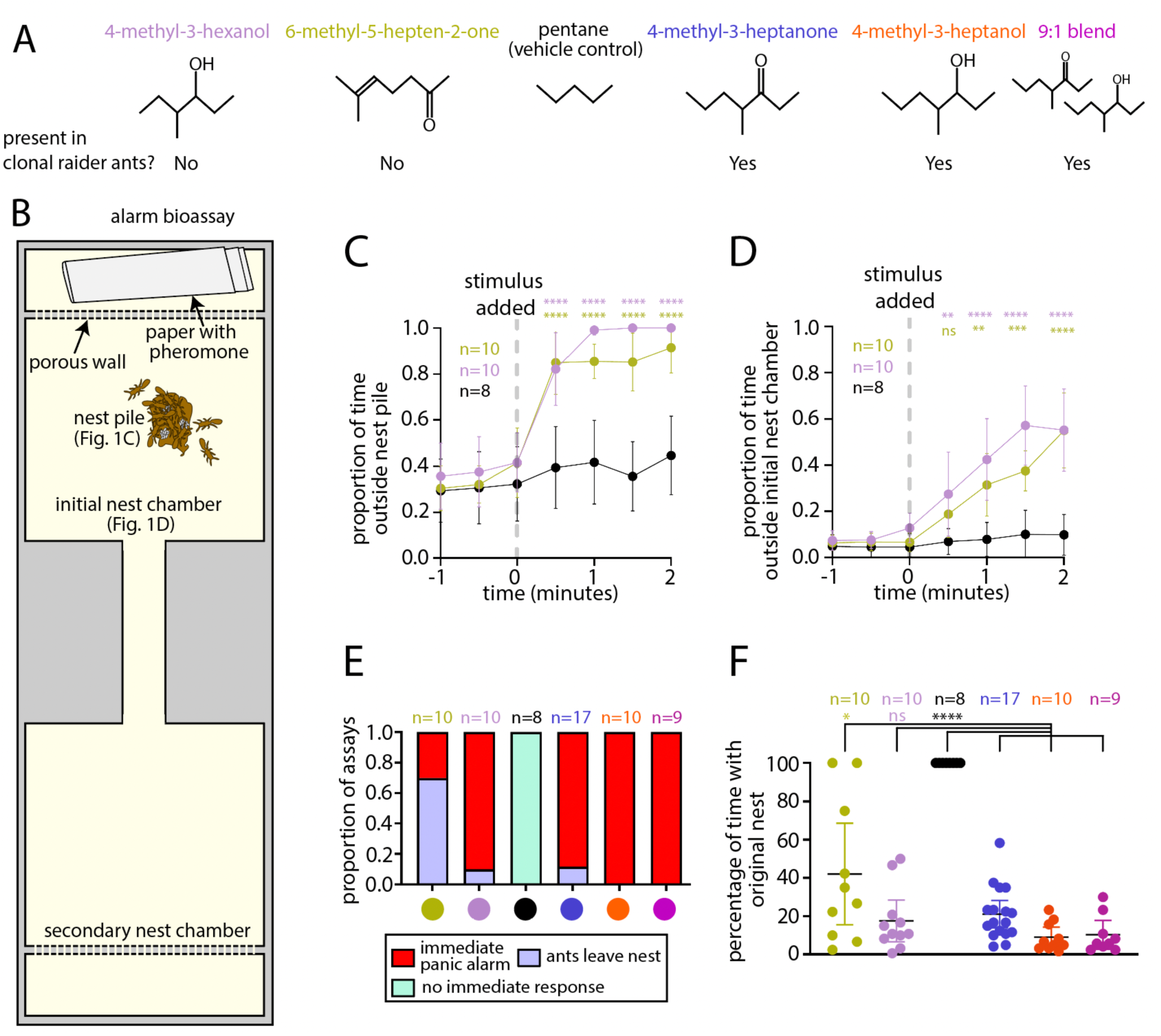
Behavioral responses to four ant alarm pheromones. (A) Chemical structures of four ant alarm pheromones and the vehicle control used in this study, obtained from the PubChem database (National Institute for Biotechnology Information: https://pubchem.ncbi.nlm.nih.gov). (B) Experimental design for the colony alarm bioassay (Lopes et al., 2022). The features used for analyses in (C-D) are indicated. (C-D) Time series of colony responses to the alarm pheromones 6-methyl-5-hepten-2-one and 4-methyl-3-hexanol vs. control, measuring the proportion of ants outside the nest pile (C) and the proportion of ants leaving the initial nest chamber (D) (mean±SEM). (E) Categorical analysis of major behavioral responses to alarm pheromone stimuli. (F) Quantification of the length of time that the original nest pile remained intact in the bioassays from (C-D); see Table S2 for details. * = p<0.05; ** = p<0.01; *** = p<0.001; **** = p<0.0001, compared to vehicle control for (C-D); non-*O. biroi* alarm pheromones and the vehicle control were compared to known *O. biroi* alarm pheromones for (F); see Table S2 for details. The color code for chemical compounds in (A) applies to all figure panels.

**Table 1.**
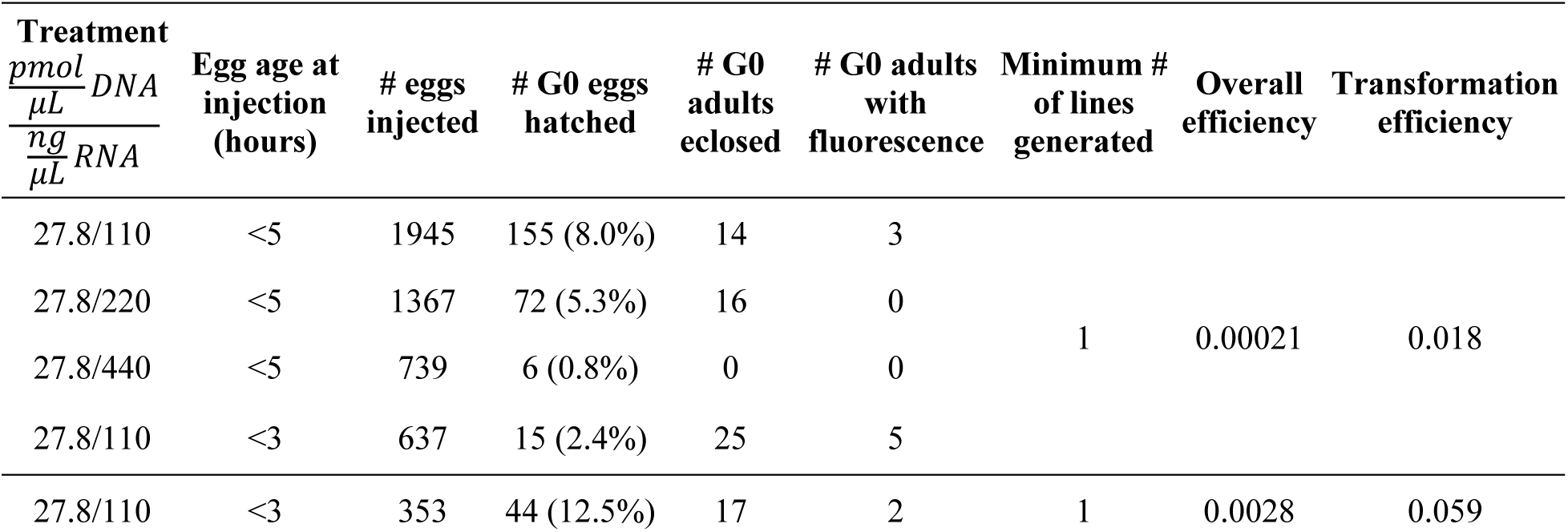
Generation of transgenic clonal raider ants expressing GCaMP6s. Injections were performed with plasmid pBAC-ie1-DsRed-ObirOrco-QF2-15xQUAS-GCaMP6s. The “Treatment” column indicates the concentrations of plasmid DNA and transposase mRNA used in the injection mix. G0 adults from the first four treatments were reared as a group, and we therefore cannot determine which treatment generated the single line that was propagated from that group. Overall efficiency was calculated by dividing the minimum number of lines generated by the number of eggs injected; transformation efficiency was calculated by dividing the minimum number of lines generated by the number of G0 adults eclosed.

Using the same bioassay and analyses that we previously used to study 4-methyl-3-heptanone and 4-methyl-3-heptanol (Fig. 1B; Lopes et al., 2022), we characterized the behavioral response to 4-methyl-3-hexanol and 6-methyl-5-hepten-2-one. Both compounds caused ants to leave the nest pile and the initial nest chamber (Fig. 1C-D). However, the behavioral responses were qualitatively distinct from one another, prompting additional analyses. Blinded categorization of the major behavioral response to each pheromone (see methods), including re-analysis of videos from our previous study (Lopes et al., 2022), showed that 4-methyl-3-heptanone, 4-methyl-3-heptanol, the 4-methyl-3-heptanone/4-methyl-3-heptanol blend, and 4-methyl-3-hexanol all caused “immediate panic alarm” in at least 80% of trials, while the most common response to 6-methyl-5-hepten-2-one was “ants leave nest”, i.e., the majority of ants slowly walking away from the nest (Fig. 1E, Supplemental Video 1).

In many of our behavioral trials, the original nest pile (defined here as the pile of eggs plus at least two workers) was disassembled, which is consistent with nest evacuation as part of a panic alarm response. In other cases, the ants moved away from the nest pile while leaving it at least partially intact, which reflects a disturbance among the ants but not a clear evacuation or panic response. We analyzed the length of time that the original nest remained intact for each odorant and found that treatment with 4-methyl-3-hexanol led to similarly rapid disassembly of the nest as 4-methyl-3-heptanone, 4-methyl-3-heptanol, and the blend (Fig. 1F, Table S2). In contrast, treatment with 6-methyl-5-hepten-2-one produced a wide range of outcomes, and the average response was significantly different from responses to clonal raider ant alarm pheromones (Fig. 1F, Table S2; Lopes et al., 2022). In summary, 4-methyl-3-hexanol elicits panic alarm behavior similarly to the native clonal raider ant alarm pheromones 4-methyl-3-heptanone and 4-methyl-3-heptanol. 6-methyl-5-hepten-2-one, on the other hand, lacks panic alarm activity and does not normally cause nest evacuation. The occasional alarm responses to 6-methyl-5-hepten-2-one could represent secondary responses, in which an ant emits actual alarm pheromone in response to the stimulus compound.

### Creation of transgenic ants

The odorant receptor co-receptor *Orco* is expressed specifically in all *O. biroi* OSNs (Trible et al., 2017; Ryba et al., 2020), and we reasoned that transgenic ants expressing GCaMP under control of an *Orco* promoter could allow optical recording of neural activity in OSN afferents in the ALs, similar to other insects (Wang et al., 2003; Stökl et al., 2010; Stensmyr et al., 2012; Zhao et al., 2022). We therefore cloned a 2.4 kb genomic fragment upstream of the *O. biroi Orco* gene which presumably contained promoter and enhancer elements sufficient to drive expression in clonal raider ant OSNs (fragment ObirOrco). We then constructed a piggyBac vector plasmid where ObirOrco drives expression of GCaMP6s (Chen et al., 2013) using the QF2 and 15xQUAS binary expression driver and effector elements in tandem to amplify transgene expression (Fig. 2A; Riabinina et al., 2015). Because we did not know if GCaMP6s would be detectable in live animals, we included an expression construct with the baculovirus-derived ie1 enhancer/promoter element to drive broad expression of the red fluorescent protein DsRed, based on similar designs used in *Drosophila melanogaster* and *Bombyx mori* (Fig. 2A; Anderson et al., 2010; Suzuki et al., 2003; Masumoto et al., 2012). We injected ant eggs with a mix of plasmid DNA and transposase mRNA (Otte et al., 2018) and reared the resulting G0 individuals using protocols modified from a previous study (see methods for details; Table 1; Trible et al., 2017). Although we generated several separate transgenic lines, we recovered a large and stable population only for the line derived from the first set of injections, and this line was therefore used for all later experiments (first four rows, Table 1). Henceforth, we refer to these ants as “GCaMP6s ants”.

**Figure 2.**
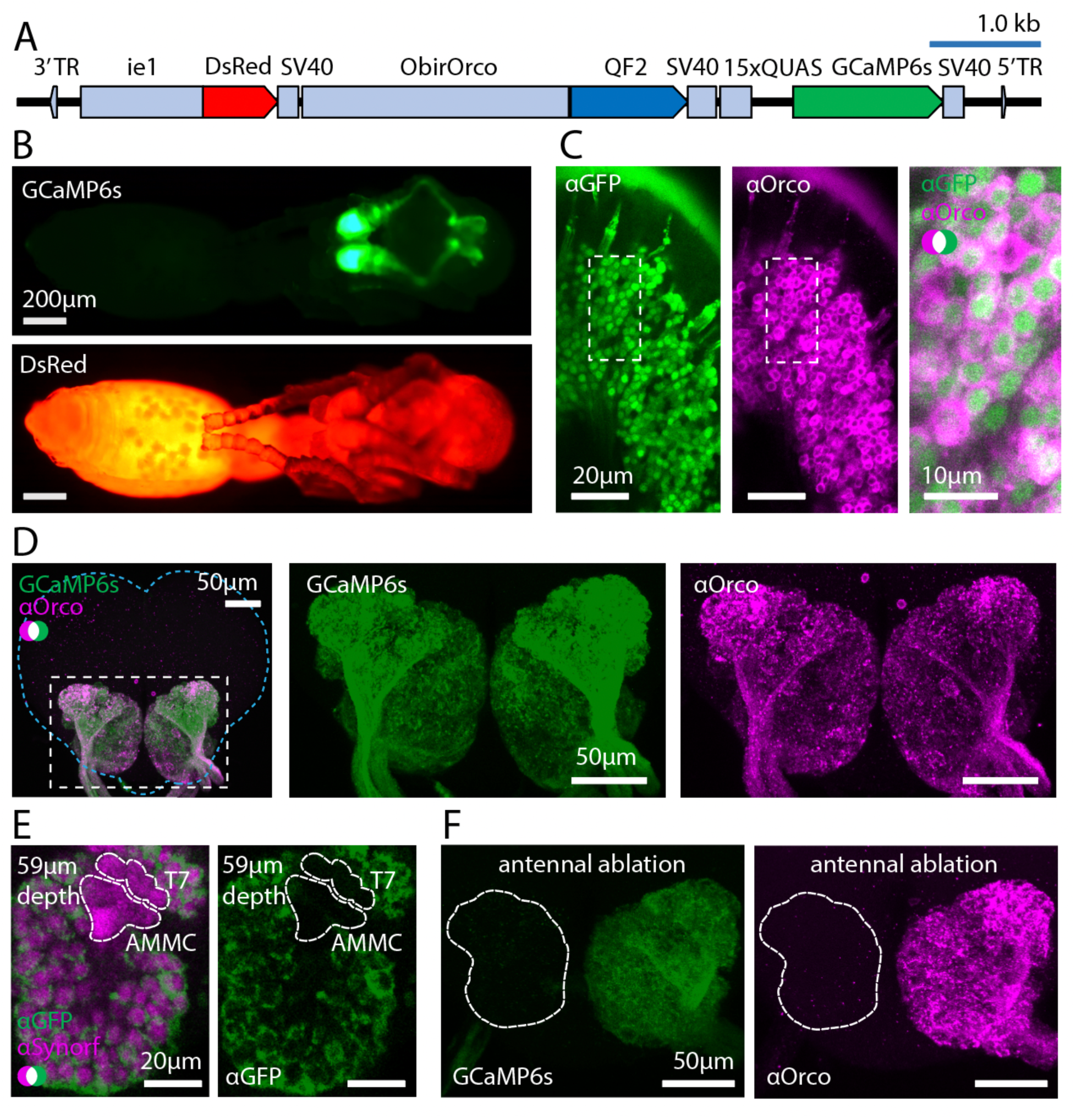
Transgene construct and GCaMP6s expression. (A) Construct design. (B) Transgene expression is easily visible in pupae viewed under epifluorescence. GCaMP6s (top); DsRed (bottom); see Fig. S1 for comparisons with autofluorescence in wild types. (C) Anti-GFP (green, cytoplasmic) and anti-Orco (magenta, membrane bound) densely label OSNs in the antennal club (max z-projection through 3 1µm slices of whole-mounted tissue). (D) GCaMP6s and anti-Orco signal co-localize in the ALs (max z-projection through the AL); brain contour is shown with cyan line. (E) All glomeruli, including the T7 cluster of the AL, are stained by anti-Synorf (neuropil; magenta). Whereas anti-GFP labeling of GCaMP6s (green) is strong in other AL glomeruli, this signal is weak or absent in the T7 glomeruli and absent in the antennal mechanosensory and motor center (AMMC). (F) Unilateral ablation of the antenna (from the scape) eliminates GCaMP6s (green, left) and anti-Orco signal (magenta, right) from the antennal lobe, indicating that GCaMP6s signal in the AL derives exclusively from sensory neurons in the antennae (max z-projections through the AL).

### Characterization of transgenic ants

We checked for GCaMP6s expression in our transgenic line to determine if it would be useful for imaging odor-evoked calcium responses. Transgenic pupae had detectable GCaMP6s fluorescence in the antennal club, consistent with expression in OSNs, and DsRed was broadly visible under epifluorescence in live animals (Fig. 2B, Fig. S1). DsRed is expressed at a low level in the AL, possibly due to leaky expression from ObirOrco (Fig. S2A). We assessed GCaMP6s expression in OSNs in the antennal club using immunohistochemistry and found that GCaMP6s labels the great majority of Orco-positive cells (Fig. 2C). Examination of brains from GCaMP6s ants showed high levels of GCaMP6s in the ALs, where it co-localizes with Orco, which labels OSN afferents (Fig. 2D). GCaMP6s is also expressed in parts of the sub-esophageal zone and central complex (Fig. S2B-C).

Examination of GCaMP6s brains stained with anti-Orco revealed that all Orco-positive glomeruli were also GCaMP6s-positive (Fig. 2D). The ~6 glomeruli of the T7 cluster are the only Orco-negative glomeruli (McKenzie et al., 2016; Ryba et al., 2020), and GCaMP6s labeling in the area mapping to the T7 cluster was weak or absent, showing specific expression of the transgene (Fig. 2E). The antennal mechanosensory and motor center (AMMC), another adjacent Orco-negative structure (Habenstein et al., 2020; Ryba et al., 2020), was also GCaMP6s-negative (Fig. 2E). Expression patterns of both DsRed and GCaMP6s were consistent across individuals. Together, this indicated that our transgenic line would in principle allow us to detect calcium responses from all olfactory glomeruli of the AL (about 99% of total glomeruli) with high specificity.

To see whether GCaMP6s is expressed by cells other than OSNs in the ALs, we performed unilateral antennal ablations on transgenic animals to sever the antennal nerve and examined their brains after allowing the fluorescent proteins to be cleared for one month. GCaMP6s and anti-Orco signals were greatly reduced across the entire AL connected to the ablated antenna, and no clear glomerular labeling remained (Fig. 2F). This indicates that GCaMP6s signal in the AL derives from the antennae and is likely to be exclusive to sensory neuron axons. GCaMP6s signal in the sub-esophageal zone and central complex was not affected by the antennal ablation (Fig. S3).

Expression of genetically encoded calcium indicators can alter cellular calcium buffering and affect behavior (Ferkey et al., 2007; Tian et al., 2012). We therefore examined whether the GCaMP6s ants had defects that could be relevant to the study of alarm pheromone sensation. We manually segmented the AL of a GCaMP6s ant and counted a total of 505 glomeruli (Fig. S4A). This is within the range of wild type ants (493-509 glomeruli; McKenzie et al., 2016; Trible et al., 2017; Ryba et al., 2020), showing that the gross AL anatomy of transgenic ants is normal. We then tested whether transgenic ants had defects in alarm behavior by subjecting GCaMP6s animals to our alarm behavior bioassay. The ants left the nest cluster in response to 4-methyl-3-heptanone, 4-methyl-3-heptanol and the blend, similar to wild types (Fig. S4B). The effect on leaving the nest chamber was only significantly different from control for 4-methyl-3-heptanone and the blend (Fig. S4C). This apparent minor difference between GCaMP6s and wild type ants could either reflect a real biological difference or result from less robust collective responses due to the smaller colony and sample sizes used in this experiment because of limited availability of GCaMP6s animals. Crucially, however, GCaMP6s ants perceive both alarm pheromones, and their behavioral response is qualitatively similar to wild types.

Finally, non-targeted transgene insertions can disrupt endogenous sequences (Bellen et al., 2011), and we therefore sequenced the genome of a GCaMP6s ant. The line contains a single, haploid transgene insertion on the 2^nd^ chromosomal scaffold (Fig. S4A-B). The insertion occurred at location Chr2:3,870,844-3,870,847, within an intron of the gene *trace amine-associated receptor 9* (Fig. S4C). Since the insertion is haploid and not within a coding region, and because GCaMP6s animals have normal AL anatomy and robust behavioral responses, these animals are well-suited for functional studies of the clonal raider ant olfactory system.

### Recording calcium responses to general odorants

We developed an *in vivo* two-photon imaging preparation for clonal raider ants, where animals are head-fixed, and a small imaging window is excised from the cuticle covering the ALs (Fig. 3A-B). Ants are then exposed to reproducible odor stimuli via a computer-controlled olfactometer (Galizia et al., 1997; Wang et al., 2003; Zube et al., 2008) and the resulting changes in GCaMP6s fluorescence are captured at 27.5fps, imaging the volume containing the entire AL every 1.2s (33 z-planes at 5µm increments; Fig. 3C-E, Supplemental Video 2). Because most clonal raider ant glomeruli are 10-20µm in diameter, they are all sampled in multiple imaging planes. Individual glomeruli were often discernible from baseline GCaMP6s fluorescence and always from calcium responses due to spatially clustered pixels with time-correlated responses (Fig. 3C). Combining volumetric imaging with a genetically encoded calcium indicator thus allowed us to record from all GCaMP6s-positive glomeruli throughout the entire AL during single odor stimulus trials, without possible confounding signals from projection neurons, lateral interneurons, or glia, and without concerns that detection of calcium responses was biased to particular AL regions (Fig. 3E).

**Figure 3.**
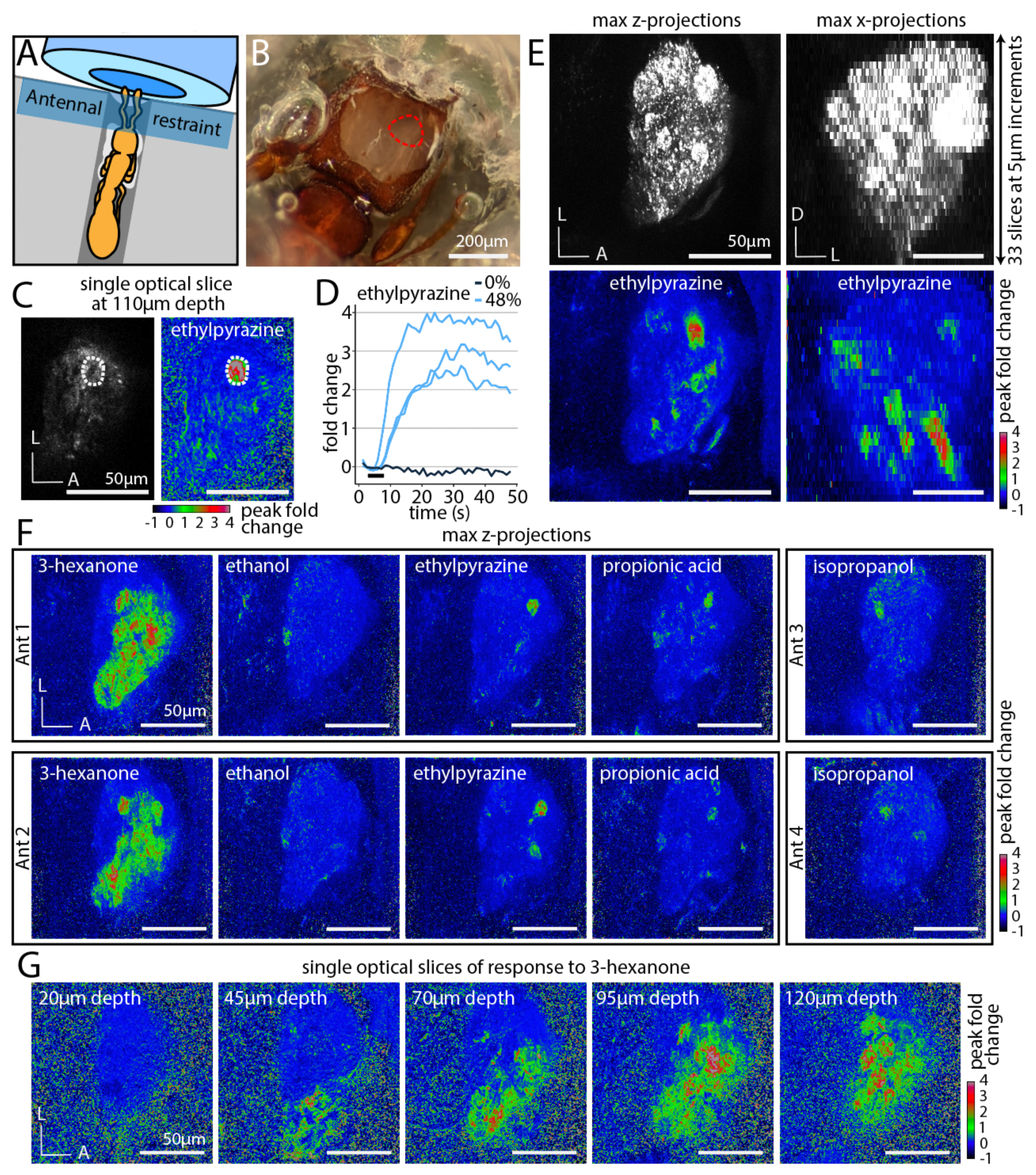
Imaging odor-evoked calcium responses in the antennal lobe. (A) A whole animal is adhered to a plastic base with glue (white) applied to the ventral side of the head and thorax. Antennae are restrained with a strip of parafilm directly in front of the air tube. (B) Preparation after dissection, with cuticle and glandular tissue removed to expose the right AL. (C) Appearance of a single optical slice through the AL using two-photon microscopy showing raw fluorescence (left, brightness and contrast enhanced) and the peak fold change of fluorescence after a 5s odor presentation at 48% concentration (right). A single glomerulus of interest is circled. (D) Time series of calcium responses in the glomerulus from (C) from several trials with ethylpyrazine or paraffin oil vehicle (0%); black bar indicates the 5s odor presentation. (E) Volumetric imaging of clonal raider ant ALs. Raw GCaMP6s fluorescence (top) is visible throughout the lobes in max z-projection (left) and max x-projection (right); responding glomeruli are visible throughout the volume (bottom) after presentation with ethylpyrazine (48%). (F) General odorants generate calcium response patterns that are qualitatively similar across different individuals (max z-projections). (G) Responses to 3-hexanone are detectable throughout the ventral/medial AL (single optical slices). D: dorsal; L: lateral; A: anterior.

To obtain a basic overview of odor representation, we presented ants (n=6) with a panel of five general (non-pheromone) volatile odorants that generated robust calcium responses in whole-AL recordings. These odorants were selected from the DoOR database of olfactory studies in Drosophila, studies of OR function in other ants, and soil volatiles (Table S3; Insam and Seewald, 2010; Münch and Galizia, 2016; Slone et al., 2017). To simplify the display of calcium responses while considering the entire AL, we calculated the peak fold change of fluorescence in each slice of the volumetric videos and then flattened them using max z-projection. Viewed this way, it was apparent that the ant AL exhibits several properties of odor encoding that have been shown in other insects (Sachse et al., 1999; Wang et al., 2003): each odorant activated a unique combination of glomeruli, and responses to the same odorant occurred in similar regions of the AL in different individuals, indicating that odor representation is qualitatively similar across individuals (Fig. 3F). We also found that the breadth of glomerular responses varied dramatically across odorants, with most odorants activating a few glomeruli, while 3-hexanone activated large regions of the ventral/medial AL (Fig. 3F-G). This demonstrates that our imaging approach can detect both sparse and broad calcium responses, if they occur.

### Pheromone representation is sparse, and alarm-inducing compounds activate a single shared glomerulus

To study encoding of alarm pheromones, we presented each ant (n=13 ants) with the four alarm pheromones at a range of concentrations (Fig. 4A). For all pheromones, max z-projections of the peak calcium response revealed sparse, unique subsets of responding AL glomeruli, while the paraffin oil vehicle did not generate responses (Fig. 4B-C). Fluorescence increases were frequently large (1-2-fold change) and lasted longer than the 5s odor presentation. We did not observe any fluorescence decreases in response to odor, although we did detect small, non-specific decreases in fluorescence due to minor shifts in AL position and photobleaching throughout the duration of each experiment. This artifact did not affect our ability to detect calcium responses, which remained robust after normalization for the duration of the experiment (Fig. S6). Comparison of calcium traces from two adjacent glomeruli showed high specificity of the response functions, without evidence for weak or transient calcium responses that might not be visible from analysis of peak fold change (Fig. S6). The response patterns to the same alarm pheromone in different individuals were qualitatively similar, in accordance with what we observed for general odorants (Fig. 3F, Fig. 4D).

**Figure 4.**
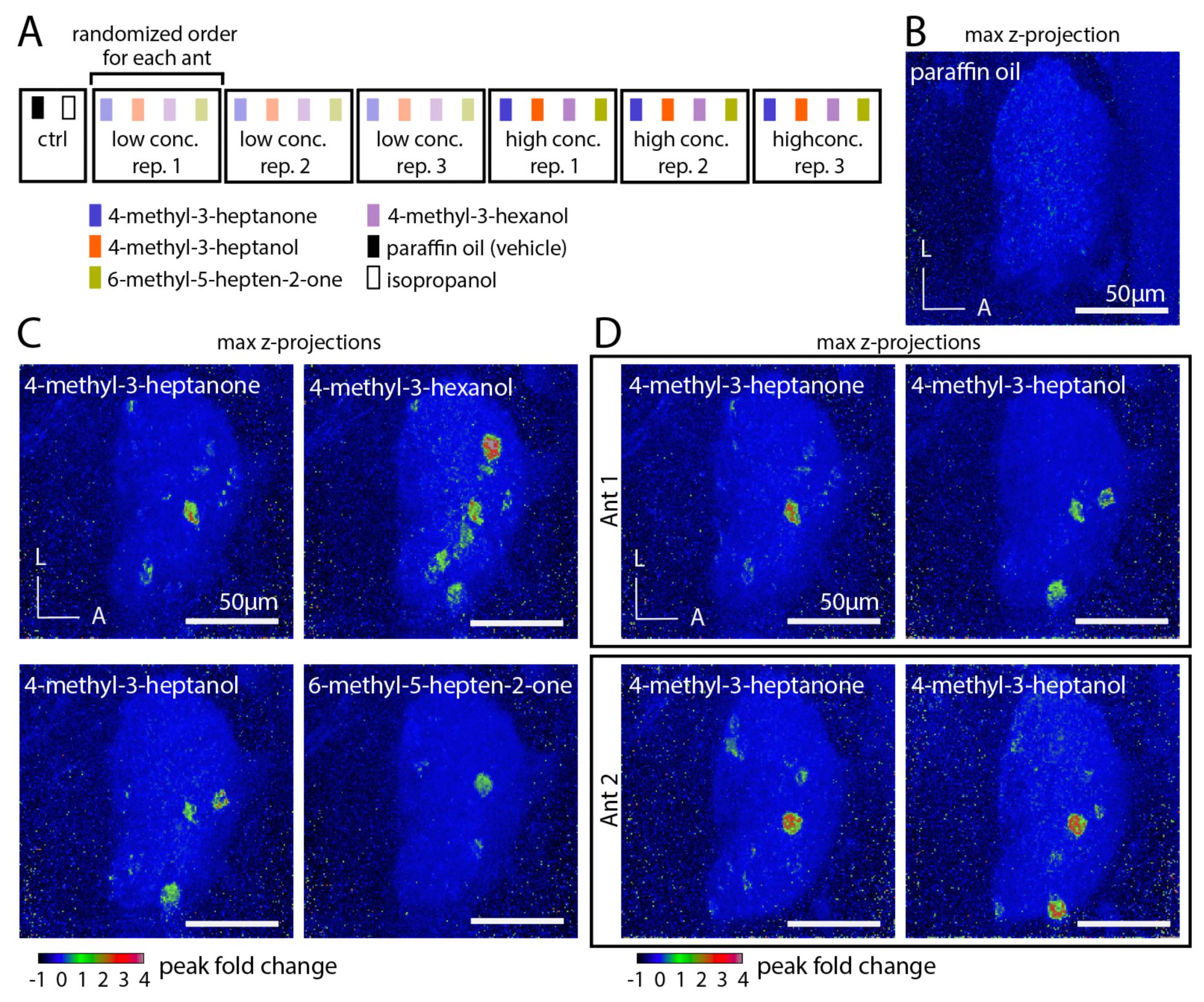
The representation of alarm pheromones in the antennal lobe. (A) Odor stimulus regime. Four alarm pheromone concentrations were tested in total (0.75%, 3.0%, 12.0%, and 48.0% v/v), but each individual ant was exposed to only two out of the four possible concentrations. (B) The paraffin oil vehicle does not generate calcium responses. (C) Representative max z-projections of peak fold change from a single ant, showing sparse activation from four alarm pheromones at high (48%) concentration. (D) Two different individuals stimulated with 4-methyl-3-heptanone (left) and 4-methyl-3-heptanol (right) at high (48%) concentration, producing qualitatively similar activation patterns. L: lateral; A: anterior.

We sought to determine how many of the ~500 glomeruli responded to each alarm pheromone by examining the max z-projections of the calcium response. We identified all regions of interest corresponding to activated glomeruli from any of the four analyzed pheromones, quantified the mean peak fold change from each pheromone/concentration, and used a threshold of ≥0.2 mean peak fold change to find robust odor-evoked responses (Fig. S7A). We observed higher numbers of responding glomeruli with increased concentration, but even at the highest concentration tested, the four pheromones activated a median of only 6 or fewer glomeruli (Fig. S7A). Despite the small number of responding glomeruli, we observed consistent partial overlap in the response patterns activated by the three compounds eliciting panic alarm responses, 4-methyl-3-heptanone, 4-methyl-3-heptanol, and 4-methyl-3-hexanol, with a single glomerulus activated by all three. We refer to this glomerulus as the “panic glomerulus, broad” (PG_b_) (Fig. S7B). This finding is consistent with the expectation that these pheromones, which can elicit slightly different forms of alarm behavior (Lopes et al., 2022), might share sensory pathways while also activating distinct sets of glomeruli. In contrast, while we sometimes observed responses to 6-methyl-5-hepten-2-one and either 4-methyl-3-heptanone or 4-methyl-3-hexanol in an overlapping region, those occurrences were rare and inconsistent (Fig. S7C).

### Alarm pheromone-responsive glomeruli are spatially stereotyped

Examining max z-projections of the calcium responses showed that PG_b_ is always located in a similar region of the AL across individuals (Fig. 5A). To better understand the level of stereotypy, we decided to localize PG_b_ more precisely, and to characterize its local environment. The raw recordings revealed that PG_b_ is located in the anterior AL, next to a region without glomeruli, approximately halfway between the dorsal and ventral AL surfaces (Fig. 5A-B). PG_b_ is neighbored by two additional glomeruli that respond to alarm pheromones, with all three visible in the same optical plane (Fig. 5B). While PG_b_ responds to 4-methyl-3-heptanone, 4-methyl-3-heptanol, and 4-methyl-3-hexanol, a nearby glomerulus responds to 6-methyl-5-hepten-2-one, which we refer to as the “6-methyl-5-hepten-2-one glomerulus” (6G). Both glomeruli were identified in 13/13 individuals. In 11/13 individuals, we identified a third neighboring glomerulus that responds to 4-methyl-3-heptanol and 4-methyl-3-hexanol, which we termed the “panic glomerulus, alcohol” (PG_a_). Examination of the position of the three glomeruli in the z-stack and comparison with a previous segmentation of the AL (McKenzie et al., 2016) showed that they are part of the T6 glomerulus cluster, which is innervated by OSNs from basiconic sensilla on the ventral surface of the ant antennal club that typically express members of the 9-exon OR subfamily (Fig. 5B; McKenzie et al., 2016). In gross anatomy, PG_b_, PG_a_, and 6G resemble typical *O. biroi* AL glomeruli and do not show obvious differences in shape or size. To validate our initial finding that these three glomeruli are functionally distinct from one another, we aligned them across individuals and quantified glomerulus-specific odor responses. This demonstrated that, while PG_b_, PG_a_, and 6G are spatially adjacent, they each reliably respond to unique combinations of odorants, with several pheromone/glomerulus combinations producing no detectable responses (Figs. 5C, S8). Importantly for its potential role in mediating alarm behavior, PG_b_ did not respond to 6-methyl-5-hepten-2-one, showing selectivity in its receptive tuning (Figs. 5B-C, S8).

**Figure 5.**
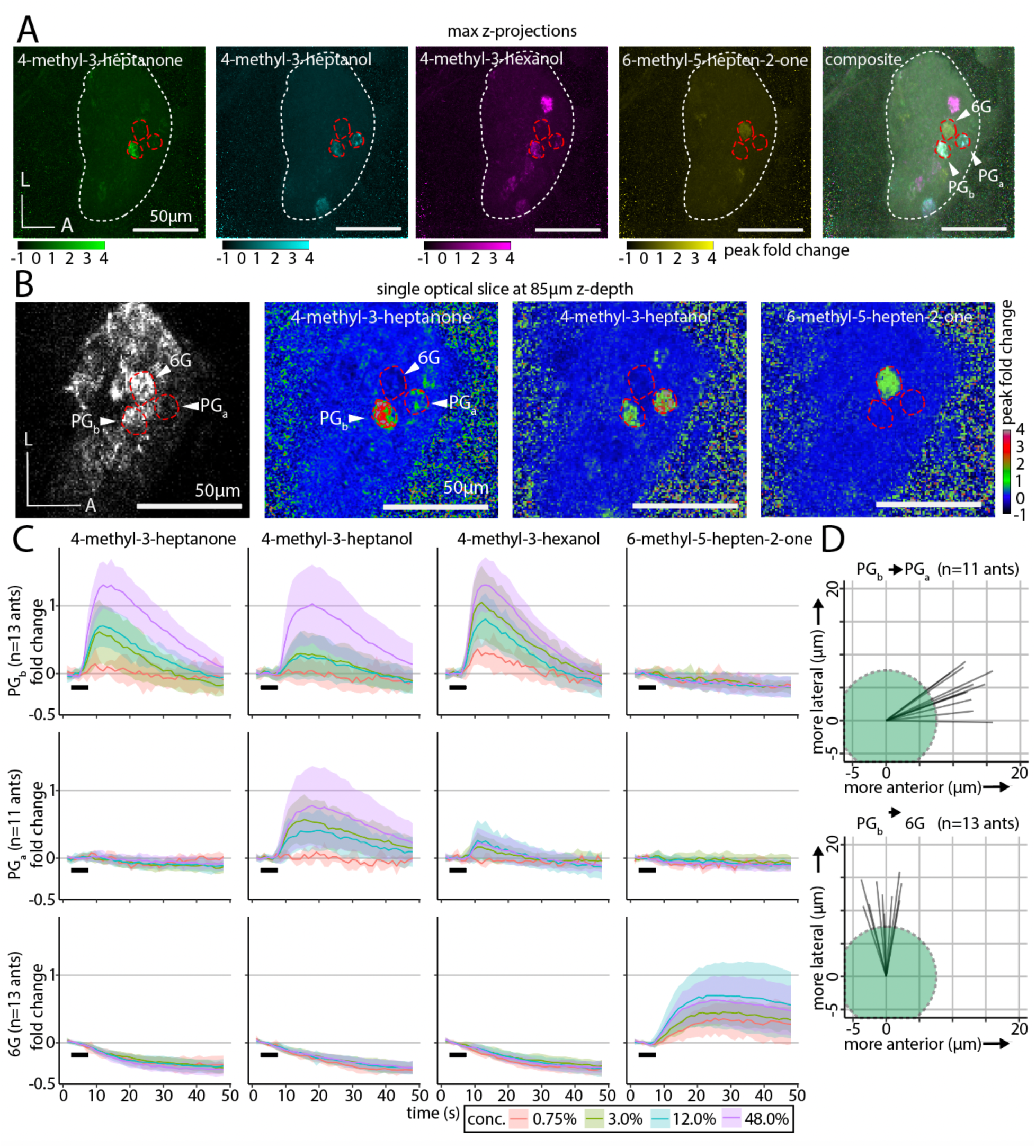
A glomerular cluster with stereotyped spatial organization and robust responses to alarm pheromones. (A) Whole-AL activation patterns for alarm pheromones overlap in several glomeruli. Three focal glomeruli are outlined. (B) Single optical slice through the AL with the three focal glomeruli (outlined). Fluorescence with enhanced brightness/contrast (left). Peak fold change in response to odors (middle left, middle right, right). See Fig. S7 for quantifications of responding glomerulus numbers at different concentrations, and Fig. S8 for peak calcium response quantifications. (C) Time series of calcium responses in PG_b_ (top), PG_a_ (middle), and 6G (bottom). Black bars indicate the 5s odor presentation. Plots show mean±SD, calculated from three trials per individual at each concentration. See Fig. S9 for extended time series of responses to 6-methyl-5-hepten-2-one in 6G. (D) Vectors of the spatial displacement between the centers of the PG_b_ and PG_a_ (top), and between the PG_b_ and 6G (bottom) glomeruli show that the spatial relationships are conserved across individuals. The green circles represent the size of a typical 15µm-diameter glomerulus, for scale. L: lateral; A: anterior.

Calcium responses had slow temporal dynamics, and in some cases calcium signals remained elevated above baseline for the duration of a single 48s recording trial. We therefore examined the temporal dynamics of alarm responses in PG_b_, PG_a_, and 6G (Fig. 5C). While responses in PG_b_ and PG_a_ had a relatively sharp peak and then declined close to baseline by the end of the 48s recording, calcium responses in 6G, which only responds to 6-methyl-5-hepten-2-one, were extremely slow, with a fluorescence plateau of tens of seconds that sometimes remained elevated at the end of the recording (Fig. 5C). We therefore performed additional odor presentations with 6-methyl-5-hepten-2-one with an extended recording period (144s) and found that calcium responses did eventually return to baseline, although in some trials the fluorescence remained elevated for >100s (Fig. S9A). At higher odor concentrations, all calcium responses lasted substantially longer than the 5s pheromone presentation (Fig. 5C). Quantifying time to response onset and time to response maximum for the different pheromones in the three focal glomeruli showed that different combinations had distinct temporal dynamics, as has been shown in other species (Fig. S9B-C; Laurent, 1999; Hallem and Carlson, 2004, 2006; Su et al., 2011).

Our analyses thus far show that alarm pheromones evoke qualitatively similar calcium responses across individuals, and that the number of activated glomeruli is consistent for a given odor. However, they do not answer the question of whether the activated glomeruli are located in fixed positions within the AL as in *Drosophila*, or whether there is significant local variation as in mice. To quantify the level of stereotypy, we examined the relative spatial positioning between PG_b_, PG_a_, and 6G along the medial-lateral and anterior-posterior axes (spatial resolution along the dorsal-ventral axis was insufficient for this analysis, especially given that these glomeruli are located at similar z-depths). We found that PG_a_ was always located anterior (mean distance between centers: 12.9±1.9SD µm), and slightly lateral (mean distance: 5.1±2.9SD µm) to PG_b_ (Fig. 5D). In comparison, 6G was always lateral to PG_b_ (mean distance: 13.1±2.6SD µm), and in a similar position along the anterior-posterior axis (mean distance: 0.6±2.2SD µm) (Fig. 5D). The standard deviation values are much smaller than the typical diameter of a glomerulus (10-20µm). We therefore conclude that these three glomeruli occupy stereotyped positions even within their local glomerular cluster and show stereotyped odor response functions across individuals.

The median number and position of responding glomeruli for each pheromone, in combination with the pheromones’ behavioral outputs, allowed us to outline a conceptual schematic of alarm pheromone representation in the ant AL (Fig. 6). The three pheromones with overlapping calcium response patterns all robustly elicited panic alarm behavior, while 6-methyl-5-hepten-2-one did not elicit panic alarm behavior and generated a non-overlapping response (Fig. 6). These findings point to a shared pathway for eliciting panic alarm behavior, centered on PG_b_.

**Figure 6.**
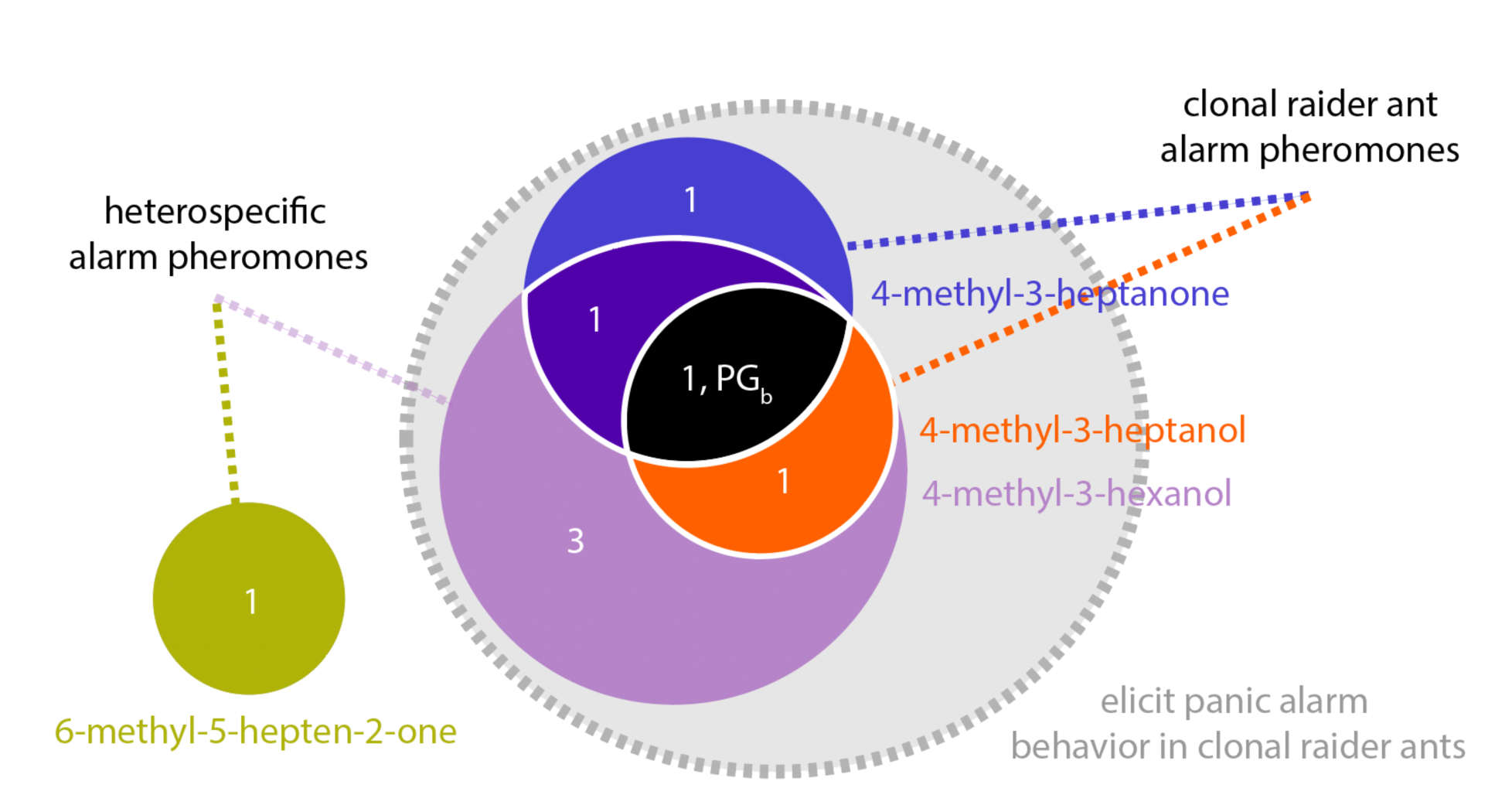
Conceptual schematic for the representation of alarm pheromones in the clonal raider ant AL. Numbers show the median number of responding glomeruli for each pheromone combination, using the highest concentration tested (48%; n=8 ants tested at this concentration). The three pheromones that elicit panic alarm responses, 4-methyl-3-heptanone, 4-methyl-3-heptanol, and 4-methyl-3-hexanol, activate mutually overlapping sets of glomeruli, while 6-methyl-5-hepten-2-one activates a mutually exclusive response in a separate glomerulus. PG_b_ is indicated on the diagram according to its response function.

## Discussion

In this study, we pioneered the combination of a genetically encoded calcium indicator with volumetric two-photon imaging to study social insect neurobiology. This allowed us to address long-standing questions about pheromone representation in the ant antennal lobe. While olfactory glomeruli in *Drosophila* occupy anatomically stereotyped positions (Stocker, 1994; Stocker et al., 1990; Gao et al., 2000; Vosshall et al., 2000; Wang et al., 2003), mouse OSNs lack hard-wired spatial targets in the olfactory bulb, with significant local variation in the spatial representation of odors across individuals (Strotmann et al., 2000; Schaefer et al., 2001). Stereotypy in the ant olfactory system, which resembles mammals in terms of complexity, has not been previously examined. We mapped a cluster of three AL glomeruli across individual clonal raider ants and found that they have consistent positions, spatial organization, and odor-evoked response functions. PG_b_, PG_a_, and 6G always had the same relative positions and occurred at similar distances from one another. Ant ALs thus possess a high degree of spatial conservation at the scale of individual glomeruli, suggesting that, similar to *Drosophila*, axon targeting by OSNs can be stereotyped, despite the vastly increased complexity of the olfactory system. However, given that our current analysis was limited to three focal glomeruli, additional work is required to determine whether this level of stereotypy is conserved across other parts of the AL.

Two of the alarm pheromones we studied are produced by the clonal raider ant, but we also investigated two additional alarm pheromones from other ant species. Of these, 4-methyl-3-hexanol elicits panic alarm behavior and activates most of the same glomeruli as the two native alarm pheromones, 4-methyl-3-heptanone and 4-methyl-3-heptanol. Thus, 4-methyl-3-hexanol represents a chemical cue emitted by other species that affects behavior (i.e., a kairomone), probably by mimicking the native pheromones via activating overlapping receptors and neural circuits. In contrast, 6-methyl-5-hepten-2-one does not robustly cause panic alarm behavior in clonal raider ants. The glomerular response pattern for 6-methyl-5-hepten-2-one is distinct from those of the panic inducing alarm pheromones, which aligns with previous work showing that compounds with different behavioral activity are usually detected through distinct olfactory channels (Isogai et al., 2011; Cattaneo et al., 2017). Interestingly, an ant-hunting spider uses 6-methyl-5-hepten-2-one as an “eavesdropping” kairomone to locate its prey, the meat ant *Iridomyrmex purpureus* (Allan et al., 1996). Given that *O. biroi* is a specialized predator of other ants, our observation that this heterospecific alarm pheromone evokes a distinct pattern of behavioral and neural activity raises the possibility that *O. biroi* may also employ kairomones to detect prey.

In both mice and *Drosophila*, olfactory glomeruli with similar chemical receptive ranges are clustered into functional subdomains, a pattern that can result from the duplication and gradual divergence of ancestral chemosensory receptors and their associated glomeruli (Uchida et al., 2000; Fishilevich and Vosshall, 2005; Prieto-Godino et al., 2016). In our experiments, all four pheromones, which share structural similarities, activated combinations of spatially adjacent glomeruli, despite the sparse representation of alarm pheromones. This suggests that the ant olfactory system also tends to map proximity in chemical space to actual spatial proximity in the AL. Here we focused on glomeruli in the T6 cluster, which are mostly innervated by OSNs expressing ORs in the 9-exon subfamily (McKenzie et al., 2016). This subfamily is particularly highly expanded via gene duplications and undergoes rapid evolution in ants (McKenzie et al. 2016; McKenzie & Kronauer 2018). Accordingly, many 9-exon ORs show similar chemical tuning (Pask et al., 2017; Slone et al., 2017). Our results are thus consistent with a model in which recently duplicated ORs are not only activated by chemically related compounds but are expressed in OSNs innervating adjacent AL glomeruli. An electrophysiological study of subsets of randomly selected olfactory projection neurons in carpenter ants also found spatially clustered responses. However, these responses came from two chemically distinct alarm pheromone components, suggesting that spatial patterning in the ant AL may also reflect pheromone social functions in addition to chemical similarity (Yamagata et al., 2006).

The proportion of glomeruli that robustly responded to any alarm pheromone was very small, with a maximum of only 6 glomeruli displaying robust activation out of ~500 total. Contrary to previous studies on social insects (Joerges et al., 1997; Galizia et al., 1998; Galizia et al., 1999; Sachse et al., 1999; Guerrieri et al., 2005; Zube et al., 2008; Brandstaetter et al., 2011; Haase et al., 2011; Carcaud et al., 2015; Paoli and Galizia, 2021), this sparse activation shows that alarm pheromones are in fact encoded by small numbers of glomeruli, similar to ecologically relevant chemicals in *Drosophila* and moths, such as sex pheromones and aversive compounds including CO_2_ and the microbial odorant geosmin (Christensen and Hildebrand, 1987; Hildebrand and Shepherd, 1997; Sakurai et al., 2004; Dweck et al., 2007; Kurtovic et al., 2007; Jones et al., 2007; Stensmyr et al., 2012). This sparse encoding logic could be advantageous by reducing the computational task for responding to molecules indicative of danger, despite the complex olfactory environment of an ant colony. With the exception of 3-hexanone, the general odorants that we tested also only activated small numbers of glomeruli. This finding is consistent with narrow tuning of individual ant ORs (Pask et al., 2017; Slone et al., 2017), and suggests that ant olfactory systems might compensate for the greater potential signal complexity implied by an expanded olfactory system by narrowing the tuning of glomeruli. Using sparse encoding for sensory signals could decrease the probability of odor mixtures activating hundreds of glomeruli simultaneously, reducing the need for vast numbers of neural connections for decoding highly combinatorial signals. We also found that the temporal dynamics of calcium responses differed by odor and glomerulus. These features provide additional information that olfactory systems can use to interpret sensory inputs, including mixtures of odors (Laurent, 1999; Hallem and Carlson, 2004, 2006; Su et al., 2011).

Ant pheromone communication employs diverse chemical substrates, including compound mixtures (Hölldobler, 1995; Morgan, 2009). These mixtures can be complex, as is the case for the cuticular hydrocarbon blends that serve as nest membership gestalt odors (Bonavita-Cougourdan et al., 1987). While ant ALs could in principle use broad encoding to represent such complex blends, insect olfactory systems can have an impressive capacity to reduce the complexity of ecologically relevant signal inputs. Mosquito ALs, for example, encode critical features of complex host odor mixtures using only a few glomeruli (Zhao et al., 2022). Future work should investigate whether the sparse encoding we reported here holds true for other types of chemical cues used by ants. This will help develop a general understanding of how glomerular tuning evolves in the context of chemical cues with high ecological relevance, complex chemical communication, and expanded olfactory systems.

## Acknowledgments

We thank members of the Kronauer, Ruta, and Vosshall labs at Rockefeller University for helpful advice and discussions. Ben Matthews provided samples of the hyPBase^apis^, pBAC-ECFP-15xQUAS_TATA-SV40, and pBac-DsRed-ORCO_9kbProm-QF2 plasmids. We thank Martin Beye for permission to use the hyPBase^apis^ plasmid, and Chris Potter for permission to use the pBAC-ECFP-15xQUAS_TATA-SV40 and pBac-DsRed-ORCO_9kbProm-QF2 plasmids prior to their appearance in publications. We thank Rob Harrell for sharing an alternate [ie1-DsRed] plasmid that was used in test injections. We thank Meg Younger for initial training and access to a two-photon microscope. We thank Daniel Pastor for sharing his protocol for immunohistochemistry in the whole-mounted antennal club. DNA sequencing was performed at the Rockefeller University Genomics Resource Center. Some confocal microscopy was performed at the Rockefeller University Bio-Imaging Resource Center. This work was supported by the National Institute of General Medical Sciences of the National Institutes of Health under Award Number R35GM127007, as well as the National Institute of Neurological Disorders and Stroke under Award Number R01NS123899 to D.J.C.K. The content is solely the responsibility of the authors and does not necessarily represent the official views of the National Institutes of Health. Additional support was provided by a Faculty Scholars Award from the Howard Hughes Medical Institute to D.J.C.K. T.H. was supported by an NSF Graduate Research Fellowship under award number 1946429 and a project grant from the Kavli Neural Systems Institute at The Rockefeller University. D.F. is an Open Philanthropy Fellow of the Life Sciences Research Foundation. L.E.L. was supported by an NSF Graduate Research Fellowship under award number DGE 194642. This work was supported in part by a grant to The Rockefeller University from the Howard Hughes Medical Institute through the James H. Gilliam Fellowships for Advanced Study program (to L.E.L. and D.J.C.K.). D.J.C.K. is an investigator of the Howard Hughes Medical Institute. This is Clonal Raider Ant Project paper number 28.

## Author contributions

T.H., W.T., L.O.C., and D.J.C.K designed the transgenics experiments. L.O.C., S.V.R., A.R., and T.H. maintained and reared ant colonies and collected eggs for injections. T.H. cloned the transgene constructs and performed the injections. L.E.L. performed and analyzed the behavioral experiments. D.F. performed the immunohistochemistry and confocal imaging and built the olfactometer. T.H. performed the epifluorescence imaging and antennal lobe reconstruction. K.D.L. prepared libraries for sequencing and performed the genomic analyses. T.H., D.F., and D.J.C.K. designed the functional imaging experiments. T.H. performed and analyzed the functional imaging experiments. T.H. and D.J.C.K. wrote the paper. All authors read, edited, and approved the manuscript for publication.

## Materials and Methods

### Behavior

#### Alarm pheromones

We purchased 96% 4-methyl-3-heptanone from Pfaltz and Bauer (Item #: M19160), and ≥99% 4-methyl-3-heptanol and 99% 6-methyl-5-hepten-2-one from Sigma-Aldrich (Item numbers M48309 and M48805-100ML, respectively). 95% 4-methyl-3-hexanol was purchased from Enamine (CAS# 615-29-2), and paraffin oil from Hampton Research (cat. #HR3-421). We also initially tested the compound undecane, which functions as an alarm pheromone in several other ant species and is found in clonal raider ant extracts (Regnier and Wilson, 1968, 1969; Ayre and Blum, 1971; Lenz et al., 2013; Lopes et al., 2022). However, undecane has a lower volatility / vapor pressure than the other alarm pheromones (Table S3), and only elicited non-specific walking behavior and no robust calcium responses in our experimental paradigms. We therefore did not investigate undecane further.

#### General odorants

98% 3-hexanone was purchased from Aldrich Chemistry (Item number 103020-10G). 98% ethylpyrazine and 99% propionic acid were purchased from Sigma-Aldrich (Item numbers 250384-5G and W292419-SAMPLE-K, respectively). 100% ethanol was purchased from Decon Laboratories (Item #: 2716), and ≥99.5% isopropanol from Fisher Chemical (Item #: A416SK-4). We initially also tested six additional general odorants with lower volatility / vapor pressure (Table S3). However, these odorants did not elicit robust calcium responses in our experimental paradigm and were therefore not studied further.

#### Colony alarm bioassay

Alarm behavior assays were performed as described previously (Lopes et al., 2022). For experiments with 4-methyl-3-hexanol and 6-methyl-5-hepten-2-one, 30 mixed-age ants from clonal line B were introduced without brood into each arena. Trials were also performed with undecane, which only induced non-specific walking behavior. For behavioral experiments with GCaMP6s ants, due to limited numbers, 15-20 ants were introduced into each arena. Prior to behavioral experiments, ants were allowed to settle for at least 5 days, until they had laid eggs and spent most of their time within a tightly packed nest pile.

Each compound (pure compounds for 4-methyl-3-heptanone, 4-methyl-3-heptanol, 4-methyl-3-hexanol, 6-methyl-5-hepten-2-one, or a 9:1 4-methyl-3-heptanone:4-methyl-3-heptanol blend) was freshly diluted 1:20 with 100% pentane each day of experiments. After recording baseline activity for 4 minutes and 30 seconds, 50 µL of each compound was added to a ~1 cm^2^ piece of filter paper and allowed to evaporate for 30 seconds before folding and placing into the stimulus chamber. Behavioral responses were recorded for another 5 minutes.

Data were analyzed as described previously, scoring three metrics of interest by hand: (1) the number of ants outside the nest pile, (2) the number of ants outside the nest chamber, and (3) the number of ants touching the mesh wall. We limited statistical analyses to the time window starting 1 minute prior to adding the stimulus and 2 minutes after. To evaluate the effect of the stimulus over time, we performed a two-way repeated measures ANOVA, and to determine the effect of the stimulus at each timepoint we used Dunnett’s multiple comparisons test.

Categorical analysis of the major behavioral response to each odorant (4-methyl-3-hexanol, 6-methyl-5-hepten-2-one, and the vehicle control, plus reanalysis of responses to 4-methyl-3-heptanone, 4-methyl-3-heptanol, and the blend from experiments in a previous study (Lopes et al. 2022) was performed by visually classifying each video as one of the following in a blinded manner: During an “immediate panic alarm”, the nest pile was disassembled within the first minute of the five-minute-long stimulus exposure. For “ants leave nest”, the nest pile persisted for at least one minute but over half of the ants left the region of the nest pile within the first minute. For “no immediate response”, the nest pile persisted for five minutes and fewer than half of the ants left the nest pile within the first minute. We also identified the time when the initial nest pile disappeared after addition of the stimulus. The nest pile was defined as the area containing the eggs and at least two adult ants. We calculated the percentage of time during which the initial nest pile remained present for the first two minutes after addition of the stimulus. We evaluated the effect of the compounds on the nest pile dissipating using a one-way ANOVA and Šidák’s multiple comparisons test to compare each additional alarm pheromone to each of the two known *O. biroi* alarm pheromones (4-methyl-3-heptanone, 4-methyl-3-heptanol, and a 9:1 blend of the two compounds).

### Generation of transgenic ants

#### Cloning and plasmid assembly

We assembled plasmid pBAC-ie1-DsRed-ObirOrco-QF2-15xQUAS-GCaMP6s using multiple rounds of PCR for generating fragments, restriction digestion with gel purification for backbones, and Gibson assembly cloning (Gibson et al., 2009; Gibson et al., 2010). Following each Gibson assembly step, correct assembly was verified using restriction digests and by sequencing PCR amplicons spanning across each of the fragment boundaries.

[1] ObirOrco: A 2.4kb promoter/enhancer fragment, including intergenic sequence and the entire 5’ UTR, amplified from clonal raider ant genomic DNA, clonal line B (NCBI LOC105284785) (primers: forward, 5’-tagttgtggtttgttgttcgcacaTATGTCACGTAATCAGCTTTTGACG-3’, lowercase shows Gibson homology region; reverse 5’-gcgcttgggtggcatgttgcaTCATATGTCTGCGAGCAAATGGAACG-3’).

[2] piggyBac backbone from pBAC-ECFP-15xQUAS_TATA-SV40 (Addgene, ID #104875) (Riabinina et al., 2016), from double restriction digest with SpeI (New England Biolabs [NEB] #R3133S) and EcoRV (NEB #R0195S).

[3] ie1: An enhancer/promoter from pGL3-IE1 (a gift from Zach Adelman, Addgene ID #52894) (Anderson et al., 2010) (primers: forward 5’-ttatcgaattcctgcagcccgggggatccaACTAGTTGTTCGCCGAGCTCTTACGCGC-3’, reverse 5’-ctcggaggaggccatCCGCGGCGAACAGGTCACTTGGTTGTTCACGATCTTG-3’).

[4] DsRed from pBac-DsRed-ORCO_9kbProm-QF2 (a gift from Christopher Potter, Addgene ID #104877) (Riabinina et al., 2016) (primers: forward 5’-acctgttcgccgcggATGGCCTCCTCCGAGAA-3’, reverse 5’-ttattatatatatattttcttgttatagatGGCGCGCCCGAACACATATGCGAACAACAAACCACAACTAGAATGCAGTG-3’).

[5] QF2 from pBac-DsRed-ORCO_9kbProm-QF2 (primers: forward 5’-aaccaagtgacctgttcgggccggACATATGCAACATGCCACCCAA-3’, reverse 5’-acccagtgacacgtgaccgCGAGCGCTGGATCTAAACGAGTTTTTAAGC-3’).

[6] 15xQUAS from BAC-ECFP-15xQUAS_TATA-SV40 (a gift from Christopher Potter, Addgene ID #104875) (Riabinina et al., 2016) (primers: forward 5’-cggtcacgtgtcact-3’, reverse 5’-tgagaacccatcgaacaagcGTTTAAACAGATCTGTTAACGAATTGATC-3’).

[7] GCaMP6s from pGP-CMV-GCaMP6s (a gift from Douglas Kim & GENIE Project, Addgene ID # 40753) (Chen et al., 2013), (primers: forward 5-gggccggcctgttcgAGCGCTTGTTCGATGGGTTCTCATCATCATC-3’, reverse 5’-atatattttcttgttatagatggCGCGCCGTAGCCCTAAGATACATTGATGAGTTTG-3’)

[8] pBAC-ie1-DsRed from Gibson assembly of piggyBac backbone, ie1-A, and DsRed fragments, transformed into NEB 10-beta competent cells (item # C3019H).

[9] ie1-B from pBAC-ie1-DsRed, (primers: forward 5’-ctgcattctagttgtggtttgttgttcgcaCATATGTGTTCGCCGAGCTCTTACGCG-3’, reverse 5’-catcgaacaagcgctcgaacaggccggcccGAACAGGTCACTTGGTTGTTCAC-3’)

[10] pBAC-ie1-DsRed-ie1-GCaMP6s from Gibson assembly of pBAC-ie1-DsRed (linearized using double restriction digest with NdeI [NEB #R0111S] and AscI [NEB #R0558S]), ie1-B, and GCaMP6s.

[11] pBAC-ie1-DsRed-ie1-QF2-15xQUAS-GCaMP6s from Gibson assembly of pBAC-ie1-DsRed-ie1-GCaMP6s (linearized using double restriction digest with FseI [NEB #R0588S] and AfeI [NEB # R0652S]), QF2, and 15xQUAS.

[12] pBAC-ie1-DsRed-ObirOrco-QF2-15xQUAS-GCaMP6s from Gibson assembly of pBAC-ie1-DsRed-ie1-GCaMP6s (linearized and second ie1 copy removed using restriction digest with NdeI) and ObirOrco.

#### Preparation of injection mixes

Plasmid DNA for injection was purified using a Machery-Nagel endotoxin-free midiprep kit (item #740420.10). The final pellet was washed under RNAse-free conditions and dissolved in nuclease-free water. To remove precipitated DNA from injection mixes, the dissolved plasmid mix was spun in a microcentrifuge at top speed for 5 minutes, and the top 90% of the supernatant was recovered. This step was repeated at least 5 times to produce injectable mix with negligible precipitate, which was stored at −20°C until injection.

We generated mRNA from the hyperactive piggyBac variant hyPBase^apis^ (Otte et al., 2018). A DNA template was generated by PCR amplification of the transposase coding sequence, with addition of a T7 promoter on the forward PCR primer, then purified using Beckman Coulter RNAClean SPRI XPBeads (item #A63987). In vitro transcription was performed with the NEB HiScribe T7 Arca mRNA kit (with tailing) (item #E2060S) to produce poly(A) tailed mRNA encoding hyPBase^apis^. The mRNA was purified using RNAClean beads (using 1.5x volume of beads compared to the reaction mix) and stored in nuclease-free water at −80°C. Template and RNA were handled under RNAse-free conditions, and a sample of mRNA was examined on an Agilent Bioanalyzer to verify RNA length and confirm absence of degradation. All DNA and RNA concentrations were measured using a Thermofisher Nanodrop.

#### Egg collection, microinjection, and larval rearing

Eggs were collected as described previously (Trible et al. 2017), with a modified schedule for treatments with eggs <3 hours old. We tested the effect of injecting even younger eggs than our previous protocol which used eggs <5 hours old (Trible et al., 2017) so that hyPBase^apis^ mRNA could be translated into active transposase while embryos still had very few nuclei, potentially reducing mosaicism. For these treatments, old eggs were removed from nests from 9am-10am, and eggs for injection were collected from 11am-11:30am, 1pm-1:30pm, 3pm-3:30pm, and 5pm-5:30pm. Injections were performed from 11:30am-12:30pm, 1:30pm-2:30pm, 3:30pm-4:30pm, and 5:30pm-6:30pm. This schedule meant that the vast majority of eggs were less than 3 hours old when injected.

Microinjections were performed as described previously (Trible et al. 2017), with the following changes: On each injection day, final injection mixes were produced by thawing and combining stored aliquots of plasmid DNA and hyPBase^apis^ mRNA under RNAse-free conditions in nuclease-free water, into a final concentration of 27.8pmol/µL plasmid and the desired concentration of hyPBase^apis^. The injected plasmid had a length of 12,025bp. The final mix was spun at top speed in a microcentrifuge for 5 minutes, and the top 90% of supernatant was used for injection. The initial mix was split into 4 aliquots and kept on ice for the day. A different aliquot was used for each round of injections. On occasions where the needle clogged, the mix was spun at top speed in a microcentrifuge before loading a new needle. The injection pressure was initially set to 3600kpa but was adjusted throughout the course of injections to maintain a consistent flow of liquid into the embryos. We varied the age of eggs and the concentration of transposase mRNA in the injection mix. Higher rates of fluorescent G0s were obtained when eggs were <3 hours old rather than <5 hours old at the time of injection. Mixes with >110ng/µL mRNA concentrations produced low hatch rates and no fluorescent G0s (Table 1).

Larvae were reared as described previously (Trible et al., 2017). Briefly, G0 larvae were hatched and placed in small colonies housed in 5cm diameter Petri dishes with a moist plaster of Paris floor to be reared by adult ants from clonal line A, which we refer to as “chaperones” when we use them to rear offspring transferred from other colonies (Trible et al., 2017). Colonies were examined under an epifluorescence microscope to confirm that some larvae expressed DsRed, indicating uptake of the plasmid.

#### Rearing initial transgenic populations

G0 individuals were reared to adulthood. For cohorts of sufficient size (~20 individuals), chaperones were removed. When the number of G0s was too small to form a robust colony, they were supplemented with wild type clonal line A ants to obtain a population of ~20 individuals. One hind leg was removed from each wild type ant to reduce their egg-laying rate compared to the G0 ants in the nest. Then, the colonies were allowed to produce G1 eggs, which were usually collected twice a week. Collected eggs were transferred to a small colony of ~20 chaperones. G1 individuals were reared to adulthood in these nests and were examined for fluorescence. Different G1 individuals potentially resulted from independent transgene insertion events. To ensure that future transgenic populations were genetically homogeneous, each fluorescent G1 adult was separated soon after eclosion, and transferred to a new transgenic line-founding colony with ~19 clonal line A ants. Eggs were collected about twice a week from these nests and given to chaperones. Fluorescent adults produced from these colonies were then returned to the transgenic line-founding colony of origin. Through several cycles of this process, genetically homogenous transgenic populations were raised and non-fluorescent individuals were removed, yielding pure colonies.

### Phenotyping transgenic ants

#### Fluorescence microscopy

Confocal microscopy of antibody-stained tissue was conducted using Zen image acquisition software on a Zeiss LSM 880 and a Zeiss LSM 900 equipped with 405nm, 488nm, 561nm and 633nm laser lines. Images were obtained using either a Zeiss LD LCI Plan-Apochromat 40X / 1.2NA or a Zeiss LD LCI Plan-Apochromat 25X / 0.8NA multi-immersion objective lens depending on the tissue sample and Zeiss Immersol G immersion medium (Zeiss # 462959-9901-000). Z-projection images were produced from stacks taken at 1µm steps using ImageJ/FIJI^98^ (Schindelin et al., 2012). Two-photon fluorescence microscopy was performed using a Bruker Investigator with a Coherent Axon laser tuned to 920nm, equipped with dual GaAsP detectors, resonant scanning galvanometer, Z-piezo module for high-speed Z-positioning, PrairieView software, and an Olympus 40X 0.9NA water-immersion objective. Images of transgenic pupae (Fig. 1B) were produced on an Olympus SZX16 epifluorescent microscope equipped with an X-Cite XYLIS light source, Olympus EP50 camera, and the appropriate filter cubes.

#### Immunohistochemistry

Antibody staining of ant brains was performed as reported previously (McKenzie et al., 2016). Briefly, the brains of female ants of a single-age cohort were dissected in cold phosphate-buffered saline (PBS) and fixed in 4% paraformaldehyde for 2 hours at room temperature. For antenna staining, a small section of cuticle was mechanically separated prior to fixation to enhance access. Blocking was performed for at least 2 hours using fresh PBS containing 0.1% or 0.5% Triton X-100 and 5% donkey serum albumin. Samples were incubated with the appropriate dilution of primary antibody in fresh blocking solution on an orbital shaker table at room temperature. Following primary incubation, samples were washed and incubated with fluorescently tagged secondary antibody diluted in fresh blocking solution. The following antibodies were used: chicken anti-GFP (Abcam #ab13970), rabbit anti-RFP (Rockland #600-401-379), mouse anti-synorf (DSHB #3C11), mouse anti-orco (gift from V. Ruta), goat anti-chicken Alexa 488 (Invitrogen #A-11039), donkey anti-mouse Alexa 647 (Invitrogen #A32787), and donkey anti-rabbit Alexa 594 (Invitrogen #A32787). For some experiments, DAPI (Invitrogen #D1306) and fluorescently tagged phalloidin (Invitrogen #A34055) were included during the secondary antibody incubation step. Stained tissue was mounted in SlowFade mounting medium on silane-coated microscopy slides (VWR #63411-01) and stored at 4°C. For AL reconstruction, a confocal stack of the right AL from a GCaMP6s-positive brain stained with anti-synorf was manually segmented using the LABKIT plugin for ImageJ, at 1µm z-axis resolution (Schindelin et al., 2012; Arzt et al., 2022).

#### Genome sequencing and genomic analyses

A single GCaMP6s ant was disrupted with a Qiagen TissueLyser II, and genomic DNA was extracted using a Qiagen QIAmp DNA Micro Kit. Libraries were prepared using Nextera Flex, and paired end, 150 base pair reads were sequenced on an Illumina NovaSeq S1 Flow Cell. Raw reads were trimmed using Trimmomatic 0.36 (Bolger et al., 2014) and aligned using bwa mem (Li et al., 2013) to both the *O. biroi* reference genome (Obir_v5.4, GenBank assembly accession: GCA_003672135.1; McKenzie and Kronauer, 2018) and a linearized plasmid reference genome created by “cutting open” the plasmid sequence at an arbitrary location on the backbone, and pasting 150 bp from the end at the front of the sequence and 150 bp from the front at the end of the sequence to accommodate any reads that might align to the vicinity of the “cut”. Reads were sorted and deduplicated using Picard (http://broadinstitute.github.io/picard/), and read depth was recorded at all sites using “samtools depth -aa,” obtaining approximately 44x coverage (Li et al., 2009). To infer the read depth of well-assembled genomic regions, we obtained all heterozygous SNPs with read depth less than 2x the genome-wide median, which excluded the fewer than 0.5% of such SNPs which likely resulted from errors in genome assembly. We then randomly selected an equal number of heterozygous SNPs as the number of base pairs in the transgene insert, and calculated read depth at those sites, and separately along both the portion of the transgene insert sequence that aligned to ObirOrco and the rest of the transgene insert.

Junction reads that aligned to both the transgene insert and the *O. biroi* reference genome were identified using the Integrative Genomics Viewer (Robinson et al., 2011), and alignments were queried by each junction read name using “samtools view” (Li et al., 2009). We performed multiple sequence alignment on these junction reads from each end of the insert using CLUSTAL 2.1 in the R package ‘msa’ (Larkin et al., 2007; Bodenhofer et al., 2015) and generated consensus sequences. To obtain the sequence of the insertion site in the reference genome, the portion of the sequence that was identical to the end of the transgene insert sequence was removed from the junction read consensus sequences. BLAST (Morgulis et al., 2008) searches of the partial consensus sequence identified a position consistent with the position these junction reads had aligned to in the *O. biroi* reference genome. The insertion locus was examined in the NCBI genome data viewer (Obir_v5.4, GenBank assembly accession: GCA_003672135.1; McKenzie and Kronauer, 2018) to check for the presence of predicted gene models.

### *In vivo* calcium imaging

#### Ant husbandry and maintenance

Ants were kept at 25°C in nests constructed by lining 5cm diameter Petri dishes with plaster of Paris. Nests were kept humidified and supplied with frozen fire ant pupae as food ~3 times per week during the brood care phase. Petri dishes held 20-80 workers each. GCaMP6s ants were propagated by cross-fostering GCaMP6s eggs into colonies with clonal line A adults (Trible et al., 2017), which were then separated into isogenic GCaMP6s colonies after eclosion. Isogenic colonies can easily be assembled in this species because *O. biroi* reproduces clonally (Kronauer et al. 2012; Oxley et al. 2014). We separated transgenic animals at the G1 stage and returned all offspring of a particular G1 individual to the same nest as their parent. For live imaging experiments, stock colonies for experiments were assembled by moving cohorts of cross-fostered GCaMP6s ants that eclosed within 2 weeks of one another into fresh Petri dish nests.

#### Specimen preparation

Adult female ants were selected from stock colonies for GCaMP imaging experiments. The age of experimental ants was 55-60- and 90-104 days post eclosion for the general odorant and alarm pheromone imaging experiments, respectively. Individuals with eyespots (indicative of intercastes; Ravary and Jaisson, 2004; Teseo et al., 2014) were excluded from our imaging study. Ants for live imaging were anesthetized on ice for ~3 minutes and then fastened to a custom two-photon imaging mount using blue-light curable glue. The antennae were restrained with a thin strip of Parafilm to decrease motion artifacts. A sheet of Parafilm with a hole for the ant’s head was applied on top of the preparation, and a watertight seal was created around the border of the head using additional glue. The preparation was then bathed with fresh ant saline (127 mM NaCl, 7 mM KCl, 1.5 mM CaCl2, 0.8 mM Na_2_HPO_4_, 0.4 mM KH_2_PO_4_, 4.8 mM TES, 3.2 mM Trehalose, pH 7.0; Zube et al., 2008) and suffused for the duration of the imaging session with additional ant saline to prevent desiccation, before excising a small imaging window in the cuticle using a sterile hypodermic needle and sharp forceps. The window was positioned above the brain, and connective and glandular tissue were removed to reveal the antennal lobes. We always imaged the right antennal lobe. Care was taken to keep the antennae and antennal nerves intact. In some cases, a muscle between the ALs and near the esophagus was severed, which reduced the amount of brain motion. This was advantageous for imaging, but not always feasible due to the small distance between the ALs and slight differences in the accessibility of the muscle from ant to ant.

#### Two-photon recording

Antennal lobe volumes were recorded at 2X optical zoom and a resolution of 512×512×33 voxels (XYZ) with 5µm Z steps, resulting in a volume with dimensions of 148×148×165µm, large enough to capture calcium transients from the entire AL which has approximate dimensions of 65µm × 125µm × 150µm. As glomeruli are typically spheroid with a diameter of 10-20µm, each glomerulus was captured in many voxels in all three dimensions. Recordings were obtained at 27.5 frames per second, resulting in 0.83 volumes per second. At the beginning of each imaging experiment, we located the dorsal surface of the AL and set that as the top of the imaging volume. We could clearly detect the boundary at the ventral surface of the AL where GCaMP6s signal disappeared, indicating that we imaged all GCaMP6s-positive glomeruli. Laser power and gain were adjusted for each ant so that all glomeruli were visible, but signal was unsaturated. Because we imaged at different depths, we compensated for loss of signal through tissue by increasing the laser power at greater depth using an exponential function. We regularly re-calibrated the position of the imaging volume, laser power, and gain in case there were any changes in baseline fluorescence or brain position during the experiment.

#### Stimulus presentation

Odors were presented using a custom-built olfactometer on 600mL/min of filtered, medical-grade air regulated with a pair of digital mass flow controllers (AliCat# MC-1SLPM-D-IPC/5M). A constant ‘carrier’ air stream (200mL/min) was presented to the ant for the duration of the imaging session to reduce mechanical stimulation of the antennae resulting from air turbulence, while a ‘stimulus’ portion of the air stream (400mL/min) was diverted and perfumed before rejoining the carrier stream at a manifold immediately upstream of the imaging preparation. By default, stimulus air bypassed control and odor vials and entered the manifold directly. During stimulus presentation, the air was perfumed by triggering high-speed three-way valves (Grainger# 6JJ52) controlled by an Arduino Uno and custom MatLab scripts, which directed the air to control or odor vials. Imaging and stimulus trials were synchronized in time using Bruker PrairieView software (i.e., the same TTL signal initiated both imaging and odor stimulation). Odors were dissolved in paraffin oil vehicle to a total volume of 300µL (concentrations represent v/v in the vial), were stored in 4mL amber glass vials with PTFE/silicone septa and connected to valves and the odor manifold via sterile hypodermic needles and nylon Luer tapers. Odor vials were prepared at the beginning of each day of imaging experiments. The air stream was directed onto the ant’s antennae using flexible PVC/vinyl tubing with an internal diameter of 1.588mm (United States Plastic Corp. Item #: 54411) from a distance of approximately 1mm.

All odor presentations had a 3s lead time and lasted for 5s. Before odor presentation, we presented the ant with the paraffin oil vehicle as a negative control and confirmed the absence of fluorescence changes before continuing the experiment. For the general odorant imaging experiment, each ant was then presented with a randomized sequence of 7-9 general odorants (48.0% concentration) which was repeated for three trials. Each of the odorants in the panel was tested in 2-6 ants. Odorants: 3-hexanone, butyric acid, dodecyl acetate, ethanol, ethylpyrazine, geranyl acetate, isopropanol, linalool, propionic acid, terpineol, and (+)-valencene. Only responses to the 5 odorants that generated robust calcium responses that were consistent across ants are shown in Fig. 3. We sometimes observed calcium activity from the other odorants, but responses were weak and not reproducible across trials in different ants. For the alarm pheromone imaging experiment, we first presented each ant with the paraffin oil vehicle and then with a positive control isopropanol stimulus. We only continued experiments with animals that showed calcium responses to the positive control but not the negative control. Each ant was presented with the four alarm pheromones in a random sequence which was first repeated for three trials at the lower concentration, followed by three additional trials at the higher concentration (for a total of 24 pheromone presentations per animal). Additional trials were performed with undecane, but these trials were not analyzed further due to absence of robust calcium responses. To reduce the impact of habituation to stimulus, each ant was presented with odors at two concentrations out of four concentrations tested (n=13 ants total, 3 ants presented with 0.75% and 12.0% odor concentrations; 2 ants with 3.0% and 12.0%; 3 ants with 12.0% and 48.0%; and 5 ants with 3.0% and 48.0%). In rare cases, we observed large motion artifacts during a recording, in which case the trial was repeated. Vials and caps were reused after cleaning as follows: removal of remaining liquid, 2x wash with 100% ethanol alternating with 2x rinse in distilled water, 2x wash with 3% Alconox alternating with 2x rinse in distilled water, 2x rinse in distilled water, air dry.

#### Image processing and analysis

Image processing was done in Fiji/ImageJ (Schindelin et al., 2012). To initially characterize response to odorants, we loaded recordings, used the “Deinterleave” function to separate them into 33 slices corresponding to videos of each recording depth, ran the Image Stabilizer plugin (Li, 2008), applied the “Gaussian Blur” filter with 1-sigma, calculated F_0_ from the mean of frames 1-5 (before any calcium changes were detected) and calculated ΔF/F_0_ by subtracting and then dividing the image stack from F_0_. The peak fold change was calculated using the “Z Project” function set to average the ΔF/F_0_ from frames 9-14, when the calcium responses typically peaked. After applying a pseudocolor LUT, we examined the peak fold change at all 33 depth positions to get a sense of the organization of glomerular responses across the ALs. We determined that all responses were positive and responding glomeruli were generally well-separated in the x/y axes. We performed additional analyses using max z-projections. Z-projections were generated by running image stabilization on each imaging plane (Li, 2008), computing ΔF/F_0_, running the “Minimum” filter with 2-pixel radius to reduce noise, applying the “Z Project” function through all slices with maximum setting, and changing all values >4 to 4 or <−1 to −1 using the “changeValues” function, to equalize the LUT range. To analyze glomerular response patterns across the whole AL, we examined all max z-projection images at the highest odor concentration for each ant and drew regions of interest (ROIs) around every glomerular region that responded to any odor in at least two trials (a small number of trials were excluded due to large motion artifacts that were only apparent after generating max projections). We then quantified the peak fold change across all trials for a particular odor and concentration and designated an ROI as responding if the value was ≥0.2. In cases where two odors activated ROIs that overlapped in the max z-projection, we examined the z-stacks to determine if the responses occurred at the same z-depth and excluded overlaps if the responses occurred at different depths. For visualizing the imaging volume (Fig. 3E, top), we used the first frame of a recording, and generated max z-projections using the z-project function. For the x-projection, we used the “Re-slice” function starting from the left to re-order the pixels, and then used the max z-project function. To visualize calcium responses throughout the imaging volume (Fig. 3E, bottom), the max z-projections of calcium responses were generated as before, but because imaging noise was more apparent in the x-projections due to higher resolution in that axis compared to the z-axis, the minimum filter was set to a 3-pixel radius.

For analyses of single glomeruli, we visually identified the z-plane containing the center of the glomerulus of interest for each trial, generated a max z-projection across 3 adjacent imaging planes (to reduce the impact of brain motion in the z-axis), and then calculated ΔF/F_0_. Peak fold change was quantified by averaging the ΔF/F_0_ over frames 9-14, the time range during which most odor-evoked calcium responses peaked.

Spatial relationships between PG_b_, PG_a_, and 6G were quantified by examining a video z-plane in which all three glomeruli were visible, placing a marker at the center of each glomerulus, and calculating the vector connecting the centers, with PG_b_ at (0,0). In two individuals, the spatial relationship between PG_b_ and PG_a_ was not quantified because PG_a_ could not be identified.

#### Statistical analyses of odor responses

We analyzed the responses of the three glomeruli PG_b_, PG_a_, and 6G to different odors and concentrations. For every glomerulus/odor combination, peak fold change values from all trials were loaded into R, and a linear regression model was fit for the peak calcium response as a function of odor concentration, with a random effect for individual, using the glm function (R Core Team, 2021). Model predictions were generated and plotted with ggplot2 (Wickam, 2016) with 95% confidence intervals.

To examine temporal dynamics in the three focal glomeruli, normalized calcium response traces from each glomerulus were loaded into R (R Core Team, 2021). The first five recorded frames were used as the baseline, and calcium response onset was defined as the latency between the start of the stimulus presentation and the time point where ΔF/F_0_ exceeded the mean of the baseline + 3SD of the baseline. The time to response maximum was defined as the latency between the start of the stimulus presentation and the timepoint with the maximum value of ΔF/F_0_. Traces where ΔF/F_0_ never exceeded the mean of the baseline + 3SD of the baseline were excluded. Only glomerulus/pheromone combinations with typically robust responses were included (4-methyl-3-heptanone in PG_b_; 4-methyl-3-heptanol and 4-methyl-3-hexanol in PG_b_ and PG_a_; 6-methyl-5-hepten-2-one in 6G). Data were plotted with ggplot2 (Wickam, 2016). To test for effects of pheromone and glomerulus identity on calcium response temporal dynamics, we performed statistical analyses on the subset of data for which the same pheromone caused responses in more than one focal glomerulus, i.e., responses in PG_b_ and PG_a_ from trials with 4-methyl-3-heptanol and 4-methyl-3-hexanol. For each temporal parameter, we built a linear mixed effects model using the lme function in R (R Core Team, 2021), modeling the effects of pheromone, glomerulus, and an interaction of pheromone/glomerulus, with a random effect for trial ID nested within ant ID.

## Supplemental information titles and legends

**Supplemental Video 1. Colony alarm bioassay.** Representative videos of each of the three categories of major behavioral response. Videos show 1 minute prior to stimulus, time skips while stimuli are put in place, then 2 minutes after stimulus (filter paper + compound). The presence of the red circle on the upper right indicates that stimulus is in place. Sped up 8x.

**Supplemental Video 2. Odor-evoked calcium response in the ant antennal lobe.** Representative two-photon recording of the calcium response to 5s presentation with ethylpyrazine (48%). A single slice corresponding to 110µm z-depth was extracted from the volumetric recording, and image stabilization was applied (Li, 2008). The “ON” label indicates the 5s odor presentation period. Sped up 3x.

**Table S1.** Background information on the four ant alarm pheromones used in this study.

| Chemical name | Example ant taxa using the compound as alarm pheromone | Major behavior in *O. biroi* (Fig. 1E) | Extracted from *O. biroi*? (Lopes et al., 2022) |
| --- | --- | --- | --- |
| 4-methyl-3-heptanone | *Atta texana* (Moser et al., 1968); *Pogonomyrmex* sp. (McGurk et al., 1966); *Neoponera villosa* (Duffield and Blum, 1973); Myrmicinae subfamily (Blum and Brand, 1972); *Atta* sp. (Hughes et al., 2001); *Eciton burchellii* & *E. hamatum* (Lalor and Hughes, 2011) | Immediate panic alarm | Yes |
| 4-methyl-3-heptanol | *Pogonomyrmex barbatus* (McGurk, 1968); *Atta sexdens* (Bento et al., 2007); Dolichoderinae subfamily (Blum and Hermann, 1978); *Harpegnathos saltator* (do Nascimento et al., 1993) | Immediate panic alarm | Yes |
| 4-methyl-3-hexanol | *Tetramorium impurum* (Pasteels et al., 1980, 1981; Morgan et al., 1992) | Immediate panic alarm | No |
| 6-methyl-5-hepten-2-one | *Formica* sp. (Duffield et al., 1977); *Iridomyrmex purpureus* (Han et al., 2022); *Aenictus rotundatus* (Oldham et al., 1994); *Eciton burchellii* (Keegans et al., 1993); *Pogonomyrmex barbatus* (McGurk, 1968); *Lasius fuliginosus* (Bernardi et al., 1967) | Ants leave nest | No |

**Table S2.** Statistical comparisons of behavioral effects. Detailed statistical comparisons for the effects of alarm pheromones on the length of time that the original nest pile remained intact in the alarm behavior colony bioassay.

| Šidák's multiple comparisons test | Mean diff. | 95% CI of diff. | Adjusted P value |
| --- | --- | --- | --- |
| 4-methyl-3-hexanol (n=10) vs.<br>4-methyl-3-heptanone (n=17) | -3.56 | -23.50 to 16.50 | 1.00 (ns) |
| 4-methyl-3-hexanol (n=10) vs.<br>4-methyl-3-heptanol (n=10) | 8.49 | -14.19 to 31.15 | 0.95 (ns) |
| 4-methyl-3-hexanol (n=10) vs. blend (n=9) | 7.20 | -16.12 to 30.52 | 0.99 (ns) |
| 6-methyl-5-hepten-2-one (n=10) vs.<br>4-methyl-3-heptanone (n=17) | 20.92 | 0.24 to 41.60 | 0.046 (*) |
| 6-methyl-5-hepten-2-one (n=10) vs.<br>4-methyl-3-heptanol (n=10) | 32.98 | 9.78 to 56.18 | 0.0012 (**) |
| 6-methyl-5-hepten-2-one (n=10) vs. blend (n=9) | 31.69 | 7.86 to 55.53 | 0.0029 (**) |
| vehicle control (n=8) vs.<br>4-methyl-3-heptanone (n=17) | 78.87 | 56.63 to 101.10 | <0.0001 (****) |
| vehicle control (n=8) vs.<br>4-methyl-3-heptanol (n=10) | 90.93 | 65.32 to 115.50 | <0.0001 (****) |
| vehicle control (n=8) vs. blend (n=9) | 86.64 | 64.43 to 114.9 | <0.0001 (****) |

**Table S3.** Vapor pressures of odorant stimuli. Odors are listed according to vapor pressure in descending order. Vapor pressure values were obtained from the PubChem database (National Institute for Biotechnology Information: https://pubchem.ncbi.nlm.nih.gov). For records with missing values from PubChem, predicted values are given instead, generated from EPISuite (US EPA, 2022) and obtained from the ChemSpider database (Royal Society for Chemistry: https://www.chemspider.com).

| Odorant | Category | Generated robust AL responses? | Vapor pressure (mmHg) at RT |
| --- | --- | --- | --- |
| ethanol | General odorant | Yes | 40 |
| isopropanol | General odorant | Yes | 33 |
| 3-hexanone | General odorant | Yes | 13.9 |
| 4-methyl-3-heptanone | Ant alarm pheromone | Yes | 5.03 (predicted) |
| propionic acid | General odorant | Yes | 2.9 |
| 6-methyl-5-hepten-2-one | Ant alarm pheromone | Yes | 1.78 (predicted) |
| ethylpyrazine | General odorant | Yes | 1.67 |
| 4-methyl-3-hexanol | Ant alarm pheromone | Yes | 1.55 (predicted) |
| 4-methyl-3-heptanol | Ant alarm pheromone | Yes | 0.43 (predicted) |
| butyric acid | General odorant | No | 0.43 |
| undecane | Ant alarm pheromone | No | 0.41 |
| linalool | General odorant | No | 0.16 |
| terpineol | General odorant | No | 0.04 |
| (+)-valencene | General odorant | No | 0.033 (predicted) |
| geranyl acetate | General odorant | No | 0.02 |
| dodecyl acetate | General odorant | No | 0.00047 |

**Figure S1.**
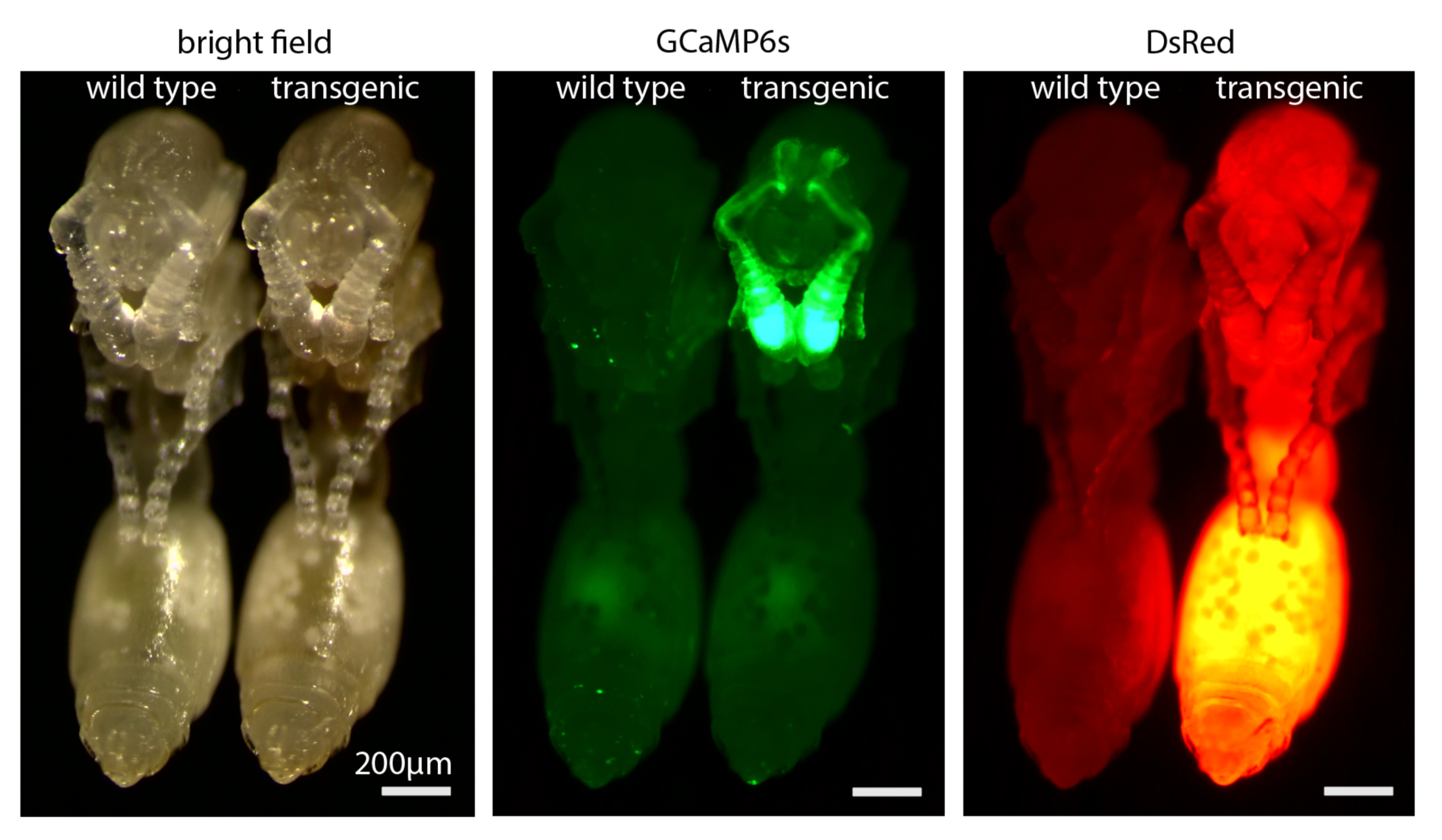
Comparison of fluorescence in transgenic vs. wild type pupae. The same two clonal line B pupae, one wild type and one transgenic, imaged under bright field (left) and epifluorescence, with filters set to detect GCaMP6s (middle) and DsRed (right). Pupae were imaged 10 days after pupation.

**Figure S2.**
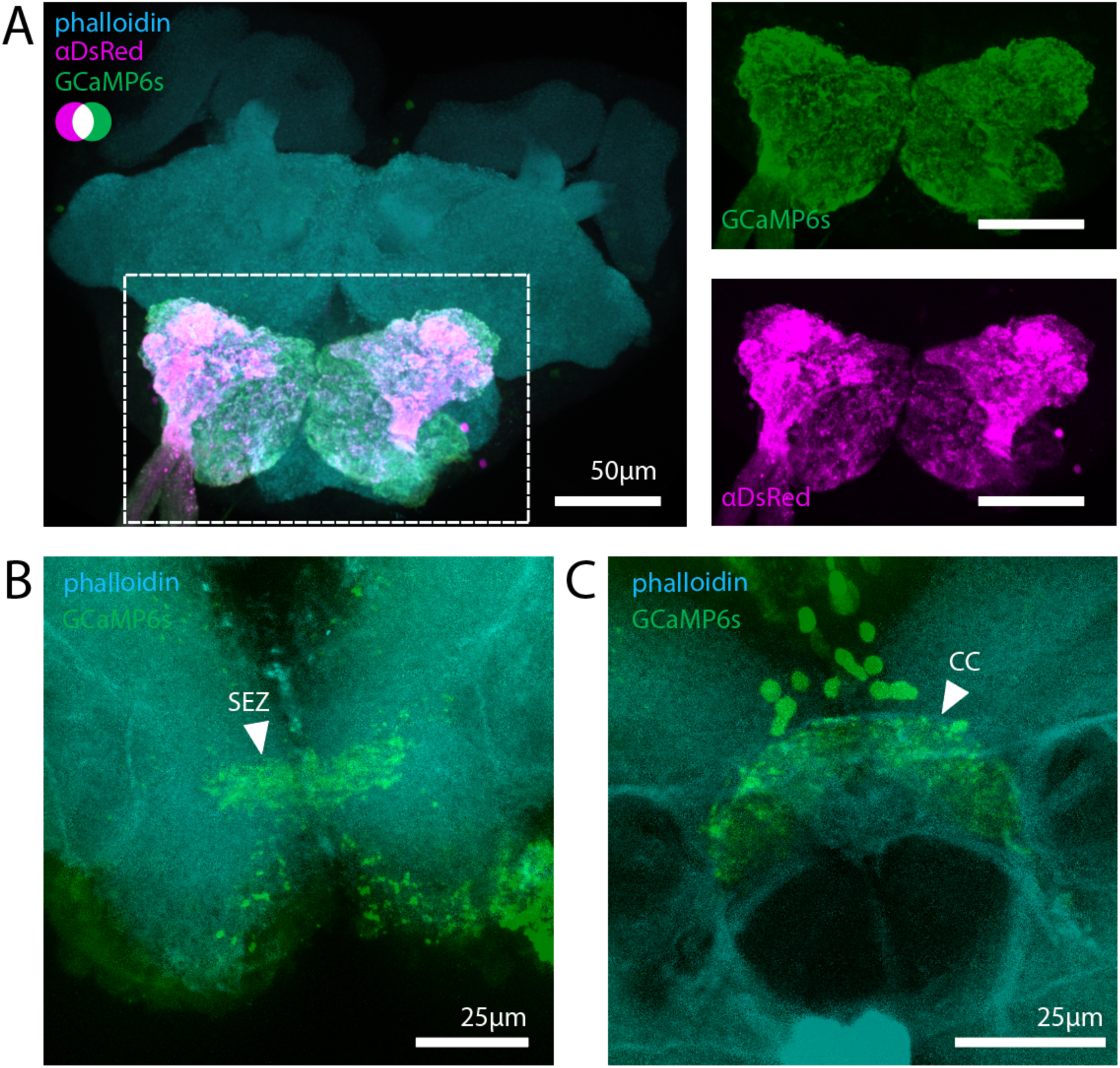
Additional characterization of transgene expression in the brain. (A) Anti-DsRed (magenta) labels the ALs, indicating co-expression of DsRed with GCaMP6s (green; endogenous fluorescence) in ants carrying [ie1-DsRed, ObirOrco-QF2, 15xQUAS-GCaMP6s]. Phalloidin stains actin (cyan). (B) GCaMP6s fluorescence (green) is detectable in the subesophageal zone (SEZ). (C) GCaMP6s fluorescence (green) is also visible in processes innervating part of the central complex (CC) as well as in a nearby cluster of somas. Images show max z-projections through the imaged brain regions.

**Figure S3.**
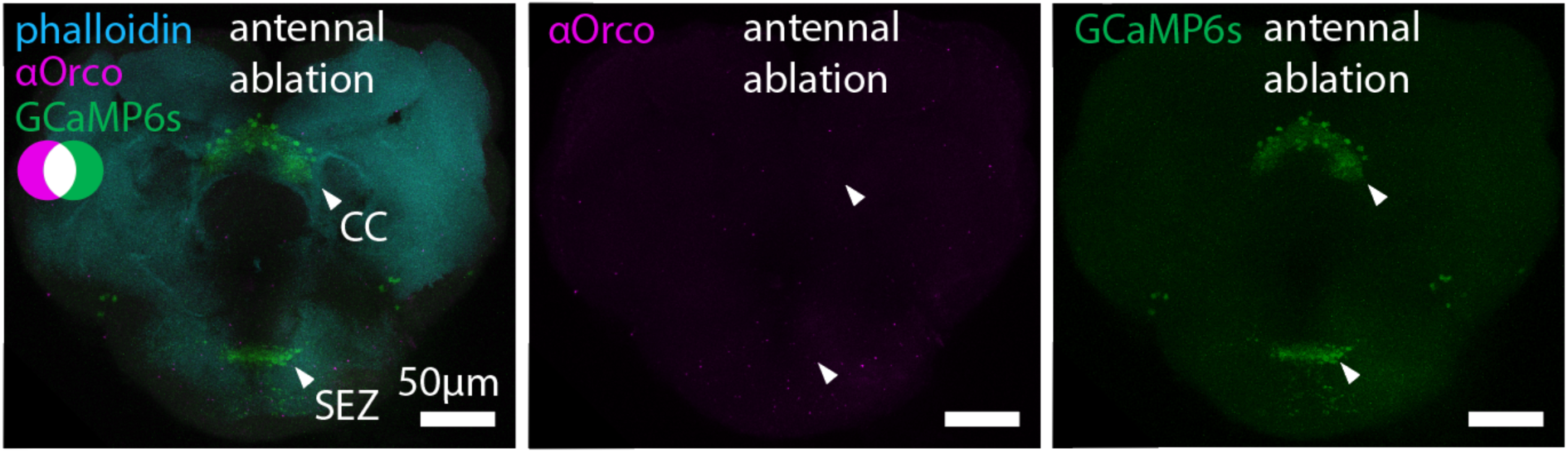
GCaMP6s signal after unilateral antennal ablation. After unilateral ablation of the antenna (from the scape), bilaterally symmetrical GCaMP6s signal is still detectable in the central complex (CC), as well as the subesophageal zone (SEZ). No anti-Orco signal was detected in these brain regions. Images show max z-projections through the imaged brain regions.

**Figure S4.**
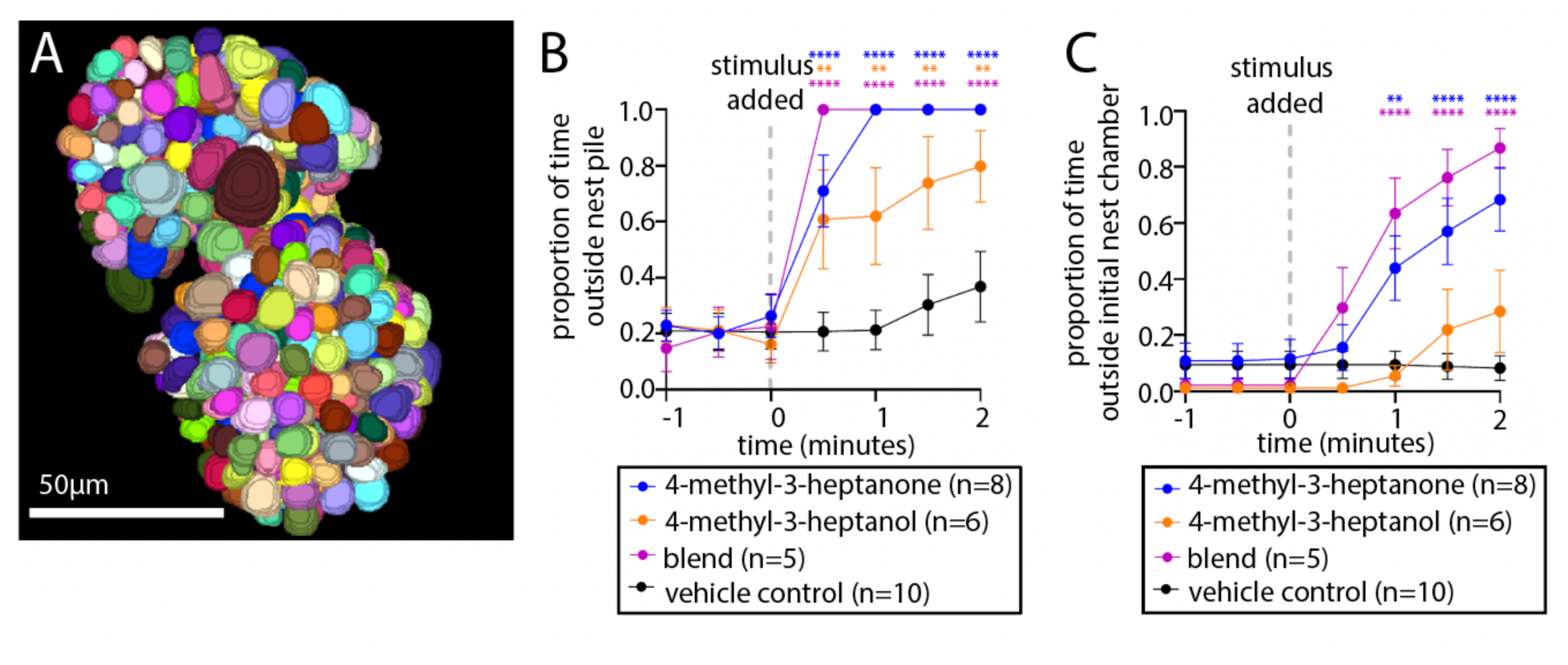
GCaMP6s ants have normal antennal lobes and respond to alarm pheromones. (A) 505 glomeruli were reconstructed from a GCaMP6s AL. (B-C) Colony alarm bioassay using GCaMP6s animals, showing mean±SEM; only comparisons that are significantly different from the vehicle control are indicated. (B) GCaMP6s animals leave the nest in response to 4-methyl-3-heptanone, 4-methyl-3-heptanol, and a 9:1 blend of the two; (C) ants leave the nest chamber in response to 4-methyl-3-heptanone and the blend. *: p<0.05; **: p<0.01; ***: p<0.001; ****: p<0.0001, compared to vehicle control.

**Figure S5.**
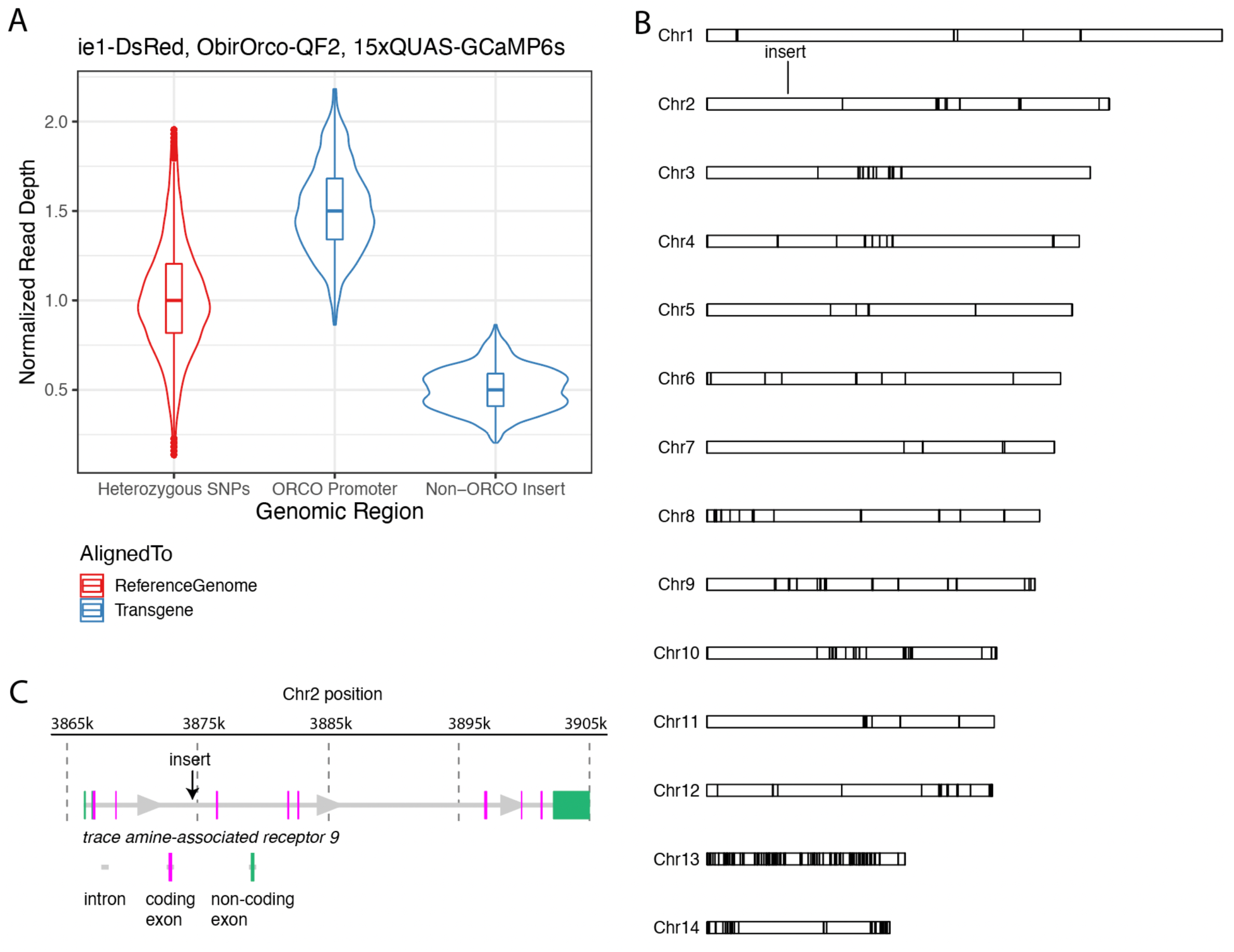
Genomic analyses of the transgenic line used for imaging. (A) Normalized read depth for reads aligning to a panel of heterozygous SNPs, the ObirOrco promoter, and the non-ObirOrco portion of the transgene. Normalized read depth for ObirOrco is ~1.5, corresponding to a single additional copy of ObirOrco inserted into the genome (added to the two endogenous copies). Normalized read depth of ~0.5 at the rest of the insert is also consistent with a single copy (haploid) insertion. (B) The transgene insert was localized to a site on the 2^nd^ chromosomal scaffold. Black bars indicate breaks between contigs. (C) Close-up of the transgene insertion locus within an intron of the gene *trace amine-associated receptor 9*.

**Figure S6.**
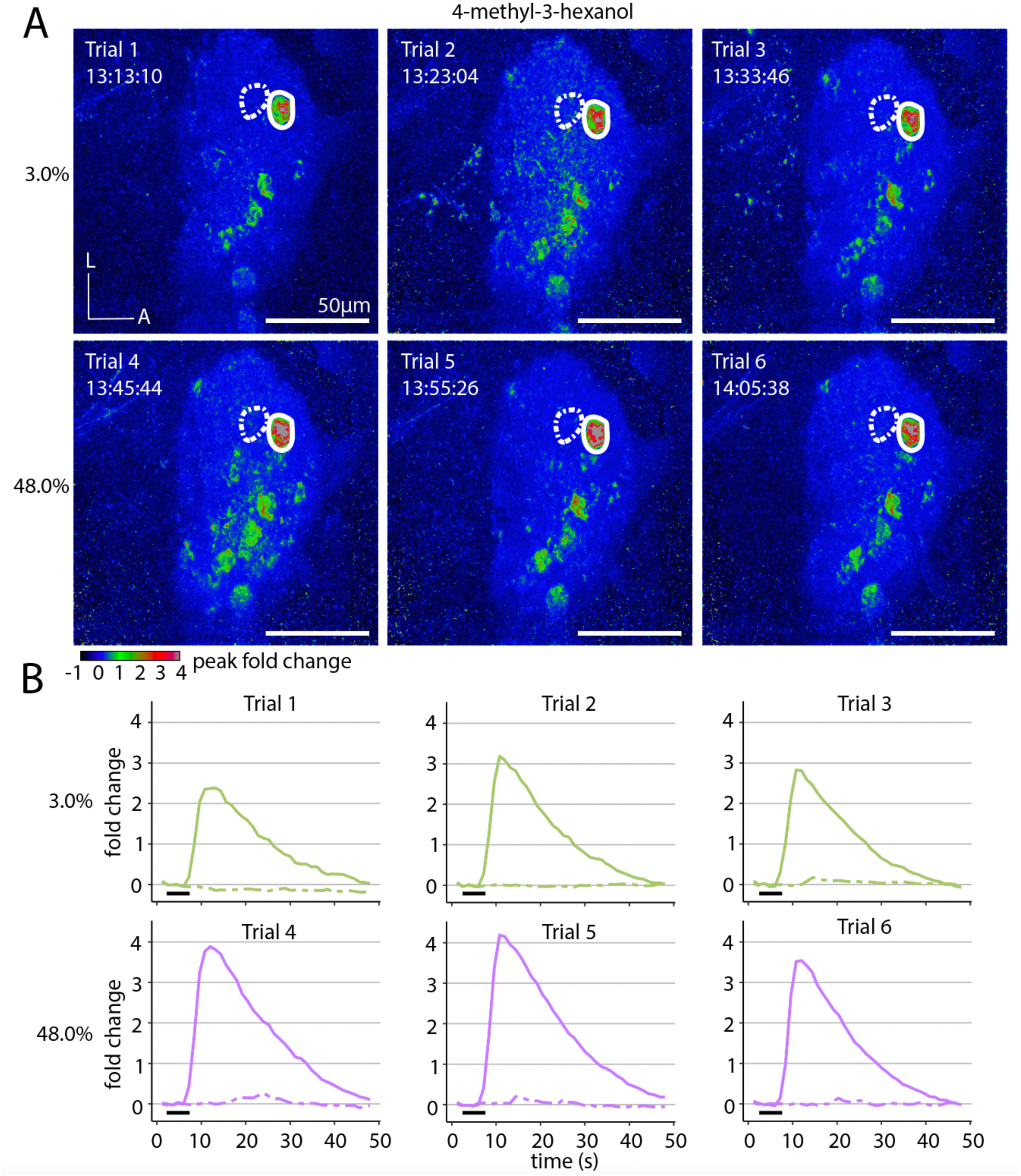
Calcium responses remain robust across trials. (A) Max z-projections of peak fold change from a single ant after presentation with 4-methyl-3-hexanol. Three trials were performed at 3.0% concentration (top), and three additional trials were performed at 48.0% concentration (bottom). Timestamps for each trial demonstrate that responses are robust over the duration of a full experiment. Two adjacent focal glomeruli are circled. (B) Time series of calcium responses from each trial in (A) for the two adjacent glomeruli; responses in the left glomerulus are shown as alternating short and long dashes and responses in the right glomerulus are shown as solid lines; black bars indicate the 5s odor presentations. Responses are quantified from max z-projections of three slices centered on 105µm z-depth.

**Figure S7.**
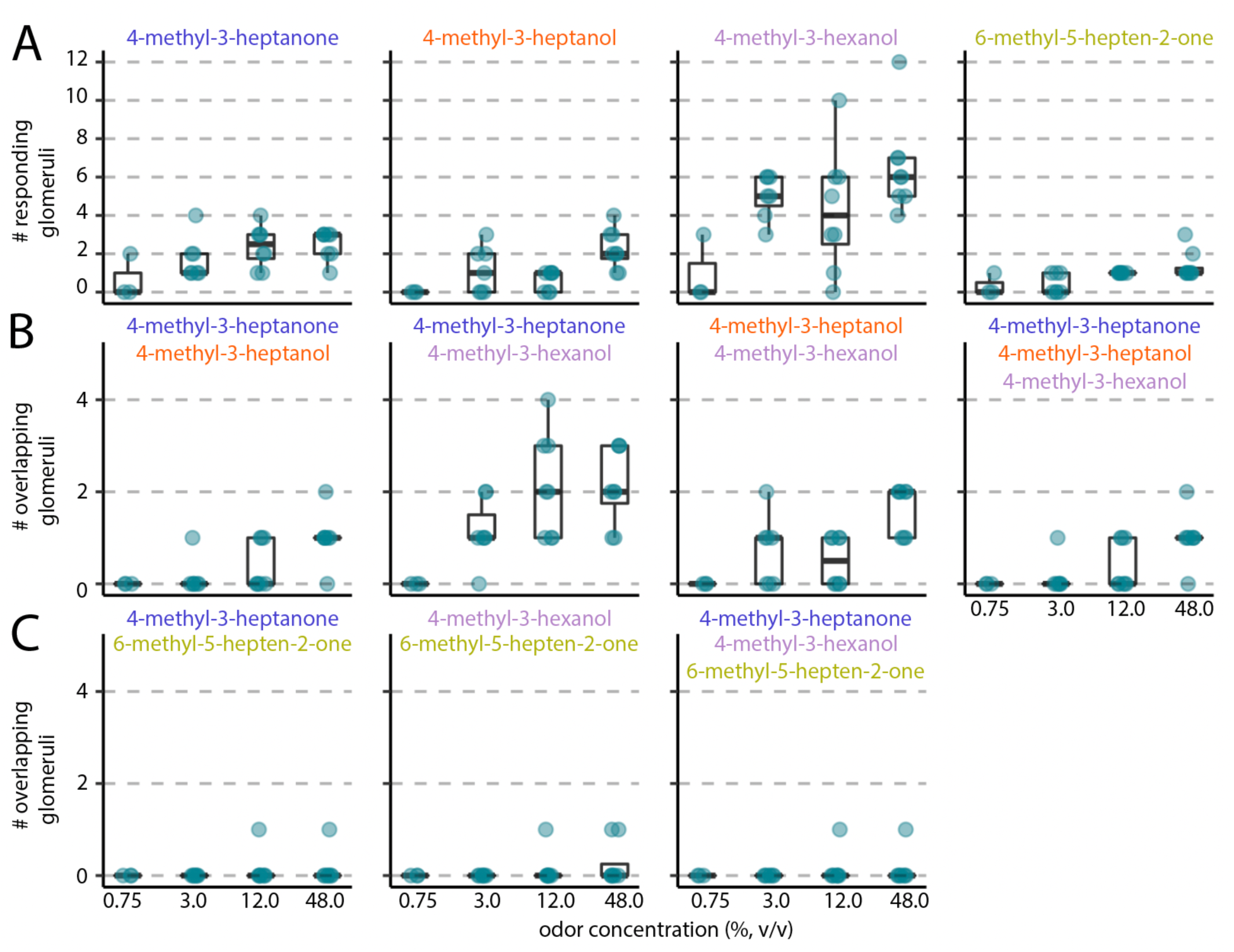
Increased odor concentration results in more responding glomeruli. Counts of the number of responding glomeruli from max z-projections; boxes enclose the first to third quartile range, with bold line showing the median and whiskers enclosing the min and max values that fall within 1.5x the interquartile range. Data points show the mean number of responding glomeruli for a given ant across all trials for a particular odorant/concentration. n=13 ants total, 3 ants presented with 0.75% and 12.0% odor concentrations; 2 ants with 3.0% and 12.0%; 3 ants with 12.0% and 48.0%; and 5 ants with 3.0% and 48.0%. Single pheromones each activated a small number of glomeruli (A), and 4-methyl-3-heptanone, 4-methyl-3-heptanol, and 4-methyl-3-hexanol activated overlapping sets of glomeruli (B). 6-methyl-5-hepten-2-one only rarely activated glomeruli shared with the other pheromones (C). All pheromones were presented separately, rather than as blends. Only pheromones and pheromone combinations that activated at least one glomerulus are shown.

**Figure S8.**
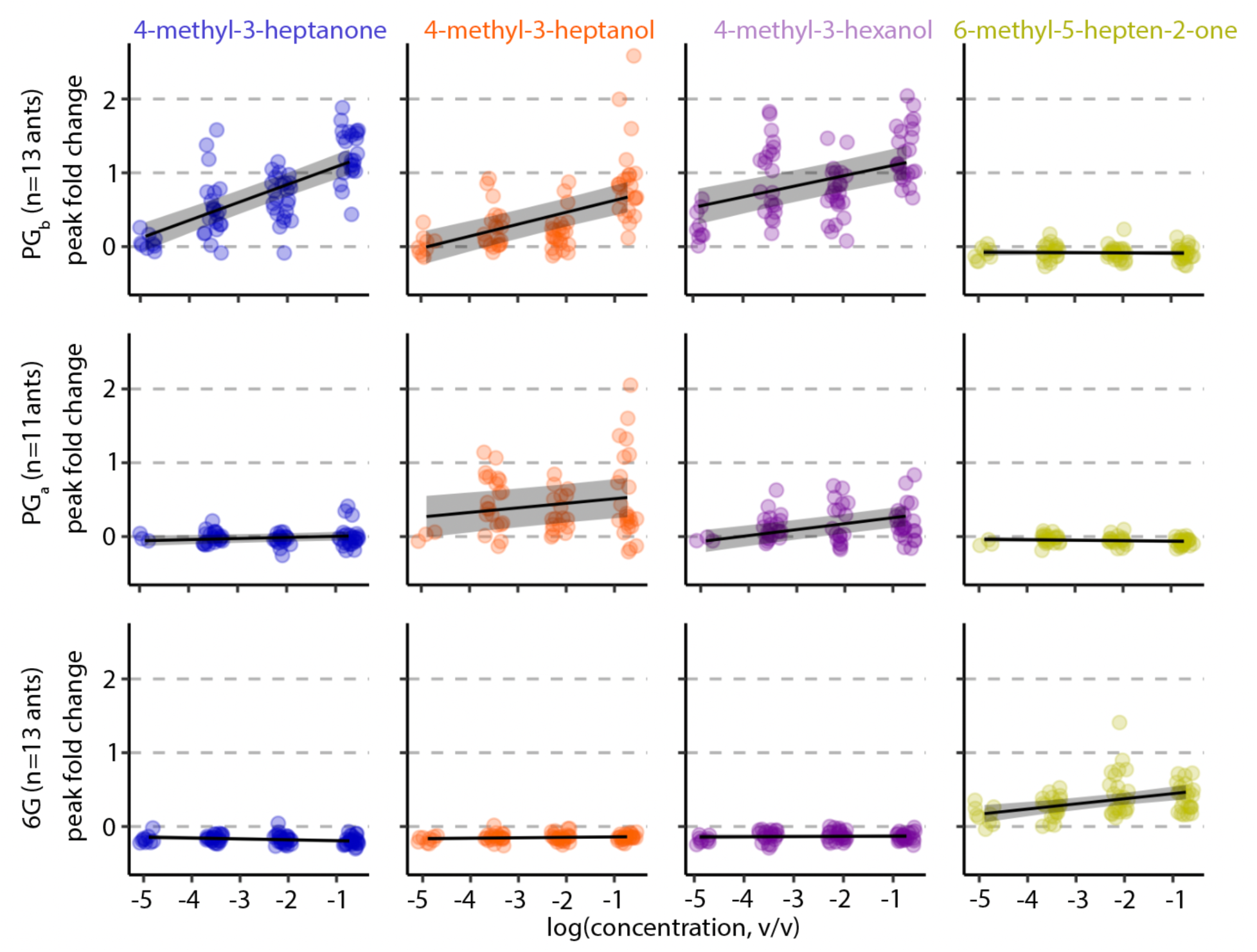
Quantification of peak fold change in PG_b_ (top), PG_a_ (middle), and 6G (bottom). Three trials per ant for each odorant/concentration; n=13 ants total, 3 ants presented with 0.75% and 12.0% odor concentrations; 2 ants with 3.0% and 12.0%; 3 ants with 12.0% and 48.0%; and 5 ants with 3.0% and 48.0%. PG_a_ could not be identified in two ants. Graphs show outputs of linear models (with 95% confidence intervals) for dose/response to each odorant in each glomerulus, with a random effect for individual. Concentrations were log transformed to show the linear relationship.

**Figure S9.**
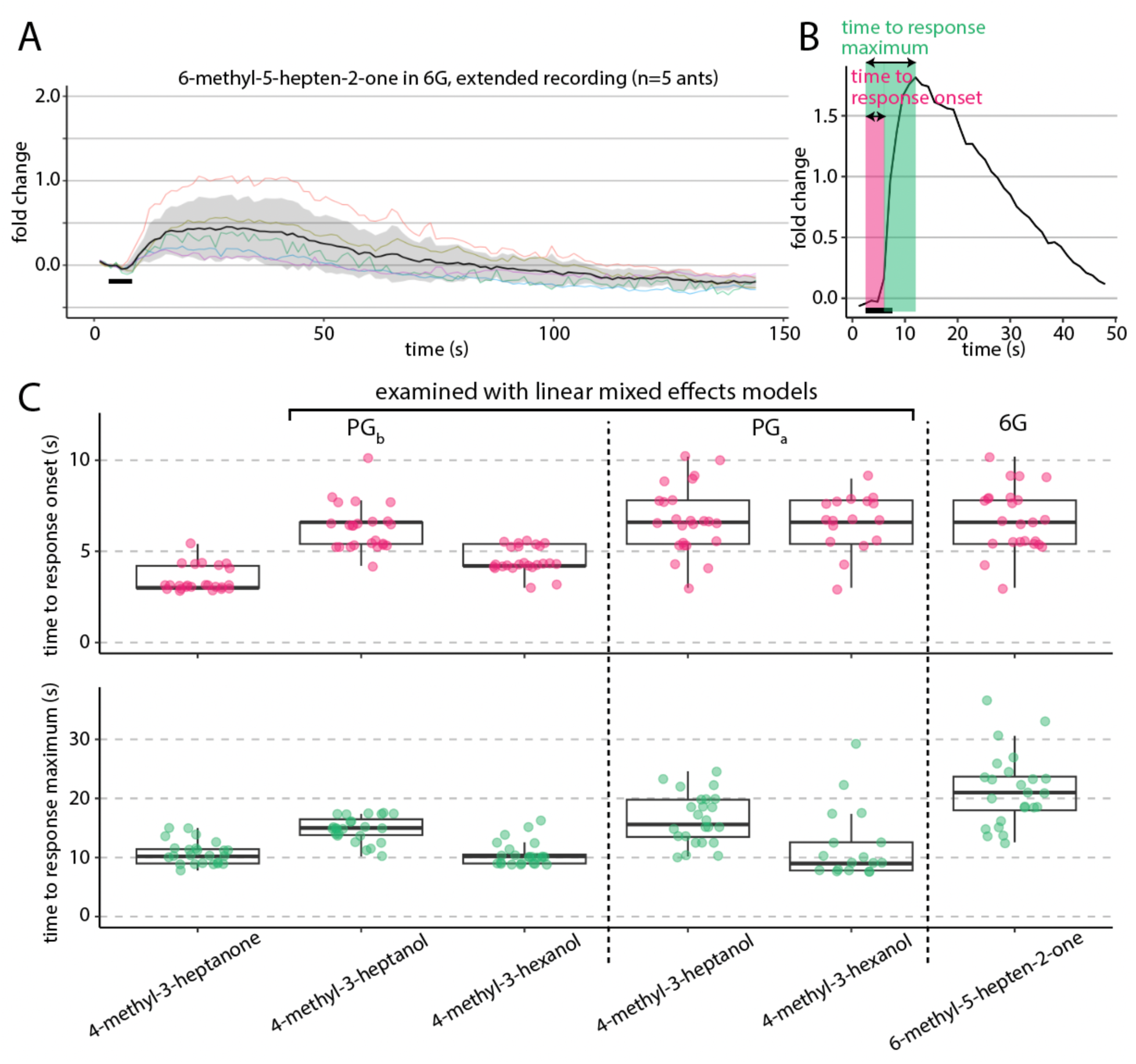
Temporal dynamics in three focal glomeruli. (A) Extended time series for the response to 6-methyl-5-hepten-2-one in 6G. Data were collected separately from Fig. 5C. Shown are single trials each from five different individuals at 48% concentration (colored traces); mean±SD (black line and gray ribbon). Fluorescence plateaued for ~30s before declining and returning to baseline ~80s after odor presentation. (B) Two parameters of temporal dynamics extracted from glomerulus-specific calcium response traces. (C) Quantification of time to response onset (top) and time to response maximum (bottom) in the three focal glomeruli PG_b_, PG_a_, and 6G in response to stimuli at 48% concentration (n=8 ants, three trials per condition per ant). Only glomerulus/pheromone combinations with typically robust responses are shown. Boxes enclose the first to third quartile range, with bold lines showing the median and whiskers enclosing the min and max values that fall within 1.5x the interquartile range. For PG_b_ and PG_a_ responses to 4-methyl-3-heptanol and 4-methyl-3-hexanol, we used linear mixed effects models to test for effects of pheromone, glomerulus, and a pheromone/glomerulus interaction on the time parameters. Time to response onset: significant effects of pheromone (p=0.0034), glomerulus (p<0.0001), and the interaction (p=0.0030). Time to response maximum: significant effects of pheromone (p<0.0001) and glomerulus (p=0.047), but not the interaction (p=0.88).

